# QuNex – An Integrative Platform for Reproducible Neuroimaging Analytics

**DOI:** 10.1101/2022.06.03.494750

**Authors:** Jie Lisa Ji, Jure Demšar, Clara Fonteneau, Zailyn Tamayo, Lining Pan, Aleksij Kraljič, Andraž Matkovič, Nina Purg, Markus Helmer, Shaun Warrington, Anderson Winkler, Valerio Zerbi, Timothy S. Coalson, Matthew F. Glasser, Michael P. Harms, Stamatios N. Sotiropoulos, John D. Murray, Alan Anticevic, Grega Repovš

## Abstract

Neuroimaging technology has experienced explosive growth and transformed the study of neural mechanisms across health and disease. However, given the diversity of sophisticated tools for handling neuroimaging data, the field faces challenges in method integration (1–3), particularly across multiple modalities and species. Specifically, researchers often have to rely on siloed approaches which limit reproducibility, with idiosyncratic data organization and limited software interoperability. To address these challenges, we have developed Quantitative Neuroimaging Environment & Toolbox (QuNex), a platform for consistent end-to-end processing and analytics. QuNex provides several novel functionalities for neuroimaging analyses, including a “turnkey” command for the reproducible deployment of custom workflows, from onboarding raw data to generating analytic features. The platform enables inter-operable integration of multi-modal, community-developed neuroimaging software through an extension framework with a software development kit (SDK) for seamless integration of community tools. Critically, it supports high-throughput, parallel processing in high-performance compute environments, either locally or in the cloud. Notably, QuNex has successfully processed over 10,000 scans across neuroimaging consortia (4), including multiple clinical datasets. Moreover, QuNex enables integration of human and non-human workflows via a cohesive translational platform. Collectively, this effort stands to significantly impact neuroimaging method integration across acquisition approaches, pipelines, datasets, computational environments, and species. Building on this platform will enable more rapid, scalable, and reproducible impact of neuroimaging technology across health and disease.

## Introduction

Neuroimaging has transformed the study of the central nervous system across species, developmental stages, and health/disease states. The impact of neuroimaging research has led to the development of a diverse and growing array of tools and pipelines that address distinct aspects of data management, preprocessing, and analysis (e.g., AFNI (5), FreeSurfer (6), FSL (7), SPM (8), HCP (9), fMRIPrep (10), QSIPrep (11), PALM (12)). However, the growing array of neuroimaging tools has created challenges for integration of such methods across modalities, species, and analytic choices. Furthermore, different neuroimaging techniques (e.g., functional magnetic resonance imaging/fMRI, diffusion magnetic resonance imaging/dMRI, arterial spin labelling/ASL, task-evoked versus resting-state etc.) have often spurred the creation of methodology-specific silos with limited interoperability across tools. This has contributed to a fragmented neuroimaging community in lieu of integrative, standardized, and reproducible workflows in the field (13).

A number of coordinated efforts have attempted to standardize acquisition and processing procedures. For example, the Human Connectome Project (HCP)’s Minimal Preprocessing Pipelines (MPP) (9) allow quality control (QC) and distortion correction for several neuroimaging modalities through a unified framework, while considering multiple formats for preserving the geometry of different brain structures (surfaces for the cortical sheet and volumes for deep structures). Another state-of-the-art preprocessing framework, fMRIPrep (10), focuses on fMRI, seeking to ensure high-quality automated preprocessing and integrated QC. QSIPrep (11) enables similar-in-spirit automated preprocessing for dMRI. FSL’s XTRACT (14) allows consistent white matter bundle tracking in human and non-human primate dMRI. Such efforts have been instrumental in guiding the field towards unified and consistent handling of data and increasing accessibility for users to state-of-the-art tools. However, these solutions are mostly application- or modality-specific, and therefore are not designed to enable an integrative workflow framework that is modality- and method-agnostic.

To address this need, we have developed the Quantitative Neuroimaging Environment & Toolbox (QuNex). QuNex is designed as an integrative platform for reproducible neuroimaging analytics. Specifically, it enables researchers to seamlessly execute data preparation, preprocessing, QC, feature generation, and statistics in a integrative and reproducible manner. The “turnkey” end-to-end execution capability allows entire study workflows, from data onboarding to analyses, to be customized and executed through a single command. Furthermore, the platform is optimized for high performance computing (HPC) or cloud-based environments to enable high-throughput parallel processing of large-scale neuroimaging datasets (such as Adolescent Brain Cognitive Development (15) or the UK Biobank (16)). In fact, QuNex has been adopted as the platform of choice for executing workflows across all Lifespan and Connectomes of Human Disease datasets by the Connectome Coordinating Facility (CCF) (4).

Critically, we have developed QuNex to integrate and facilitate the use of existing software packages, while enhancing their functionality through a rich array of novel internal features. Our platform supports a number of popular and well-validated neuroimaging tools, with a framework for extensibility and integration of additional original packages (see **Discussion**). Moreover, QuNex offers functionality for onboarding entire datasets, with compatibility for the BIDS (Brain Imaging Data Structure, (17)) or HCP-style conventions, as well as support for NIFTI (volumetric), GIFTI (surface meshes), CIFTI (grayordinates), and DICOM file formats. Lastly, QuNex enables analysis of non-human primate (18) and rodent (e.g. mouse) (19) datasets in a complementary manner to human neuroimaging workflows. To our knowledge, no existing framework provides comprehensive functionality to handle the diversity of neuroimaging workflows across species, modalities, pipelines, analytic workflows, datasets, and scanner manufacturers, while explicitly enabling methodological extensibility and innovation. Thus, QuNex provides a novel, integrative solution for consistent and customizable workflows in neuroimaging.

QuNex offers an integrative solution that minimizes technical bottlenecks and access friction for executing standardized neuroimaging workflows at scale with reproducible standards. In this paper, we present QuNex’s capabilities through specific example use cases: 1) Turnkey execution of neuroimaging workflows and versatile selection of data for high-throughput batch processing with native scheduler support; 2) Consistent and standardized processing of datasets of various sizes, modalities, study types, and quality; 3) Multimodal feature generation at different levels of resolution; 4) Comprehensive and flexible general linear modeling at the single-session level and integrated interoperability with third-party tools for group-level analytics; 5) Support for multi-species neuroimaging data, to link, unify and translate between human and non-human studies. For these use cases, we sample data from over 10,000 scan sessions that QuNex has been used to process across neuroimaging consortia, including clinical datasets.

## Results

Through QuNex, researchers can use a single platform to perform onboarding, preprocessing, QC and analyses across multiple modalities and species. We have developed an opensource environment for multi-modal neuroimaging analytics. QuNex is fully platform-agnostic and comes in the form of both Docker and Singularity containers which allows for easy deployment regardless of the underlying hardware or operating system. It has also been designed with the aim to be community-driven. To promote community participation, we have adopted modern and flexible development standards and implemented several supporting tools, including a SDK that includes helper tools for setting up a development environment and testing newly developed code, and an extensions framework through which researchers can integrate their own pipelines into the QuNex platform. These tools enable users to speed up both their development and integration of newly developed features into the core codebase. QuNex comes with an extensive documentation both in the format of inline help through the command line interface (CLI) and a dedicated Wiki page. Furthermore, users can visit our forum (https://forum.qunex.yale.edu/) for anything QuNex related, from discussions to feature requests, bug reports, issues, and usage assistance.

### QuNex is an Integrative Multi-modal and Multi-species Neuroimaging Platform

Given the diversity of sophisticated tools for handling neuroimaging data, the field faces a key challenge around method integration. We addressed this challenge by building a platform for seamless integration of a wide array of neuroimaging operations, ranging from low-level onboarding of raw data to final cutting-edge surface-based analyses and visualizations. **Figure 1** provides a general overview of the QuNex platform, while a summary of QuNex commands and functionalities is shown in **Figure S1**. QuNex supports processing of diverse data from multiple species (i.e., human, macaque and mouse), modalities (e.g., T1w, T2w, fMRI, dMRI), and common neuorimaging data formats (e.g., DICOM, NIfTI and vendor-specific Bruker and PAR/REC). It offers support for onboarding of BIDS-compliant or HCP-style datasets and native support for studies that combine neuroimaging with behavioral assessments. To this end, it allows for integrated analyses with behavioral data, such as task performance or symptom assessments, and provides a clear hierarchy for organizing data in a study with behavior and neural modalities (**Figure S2**). QuNex is capable of generating multi-modal imaging-derived features both at the single subject level and at the group level. It enables extraction of structural features from T1w and T2w data (e.g., myelin, cortical thickness, volumes, sulcal depth and curvature), white matter microstructure and structural connectivity features from dMRI data (e.g., whole-brain “dense” connectomes, regional connectivity, white matter tract segmentation) and functional features from fMRI data (e.g., activation maps and peaks, functional connectivity matrices or connectomes). As described below and shown in Figure 4, features can be extracted at the dense, parcel, or network levels using surface or volume-based analysis.

**Fig. 1.**
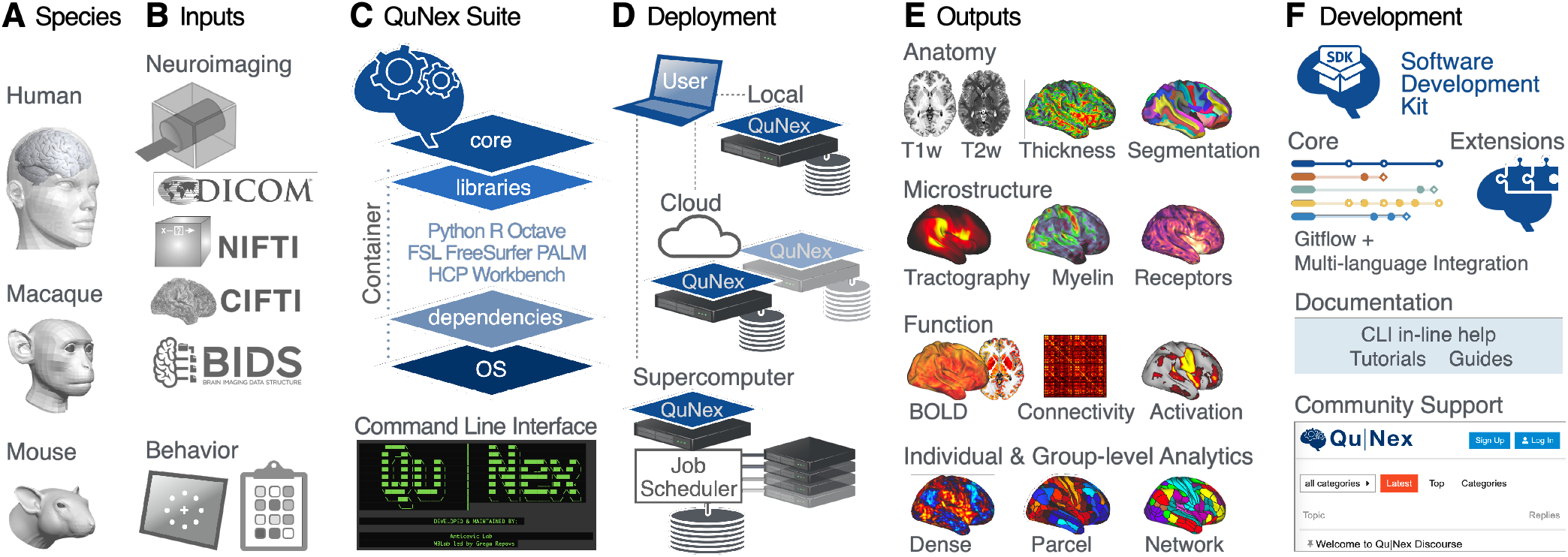
QuNex Provides an Integrated, Versatile and Flexible Neuroimaging Platform. **A)** QuNex supports processing of input data from multiple species, including human, macaque and mouse. **B)** Additionally, data can be onboarded from a variety of popular formats, including neuroimaging data in DICOM, PAR/REC, NIfTI formats, a full BIDS dataset, or behavioral data from task performance or symptom assessments. **C)** The QuNex platform is available as a container for ease of distribution, portability and execution. The QuNex container can be accessed via the command line and contains all the necessary packages, libraries and dependencies needed for running processing and analytic functions. **D)** QuNex is designed to be easily scalable to accommodate a variety of datasets and job sizes. From a user access point (i.e. the user’s local machine), QuNex can be deployed locally, on cloud servers, or via job schedulers in supercomputer environments. **E)** QuNex outputs multi-modal features at the single subject and group levels. Supported features that can be extracted from individual subjects include structural features from T1w, T2w and dMRI (such as myelin, cortical thickness, sulcal depth and curvature) and functional features from BOLD imaging (such as functional connectivity matrices). Features can be extracted at the dense, parcel, or network levels. **F)** Importantly, QuNex also provides a comprehensive set of tools for community contribution, engagement and support. A Software Development Kit (SDK) and GitFlow-powered DevOps framework is provided for community-developed extensions. A forum (https://forum.qunex.yale.edu) is available for users to engage with the QuNex developer team to ask questions, report bugs and/or provide feedback.

### Turnkey Engine Automates Processing via a Single Command

Efficient processing of neuroimaging datasets requires streamlined workflows that can execute multiple steps, with minimal manual intervention. One of the most powerful QuNex features is its “turnkey” engine, accessible through the `run_turnkey` command. The turnkey functionality allows users to chain and execute several QuNex commands using a single command line call, enabling the generation of consistent outputs in an efficient, streamlined manner. The turnkey steps are entirely configurable and modular, such that users can customize workflows to suit their specific needs. An example of an end-to-end workflow is shown in **Figure 2A**. The QuNex turnkey engine supports data onboarding of the most commonly used neuroimaging formats, state-of-the-art preprocessing pipelines (e.g., HCP MPP (9), see **Figure S3**) and denoising techniques, as well as steps for data QC. QuNex expands upon preprocessing functionalities offered by other packages by providing robust QC functionality, via visualizing key features of multi-modal data (including T1w, T2w, dMRI, and BOLD, **Figure S4**). This simplifies thorough validation of the quality of input data as well as the intermediate and final preprocessing outputs. Users can additionally choose to generate neuroimaging features for use in further analyses, including the parcellation of timeseries and functional connectivity.

**Fig. 2.**
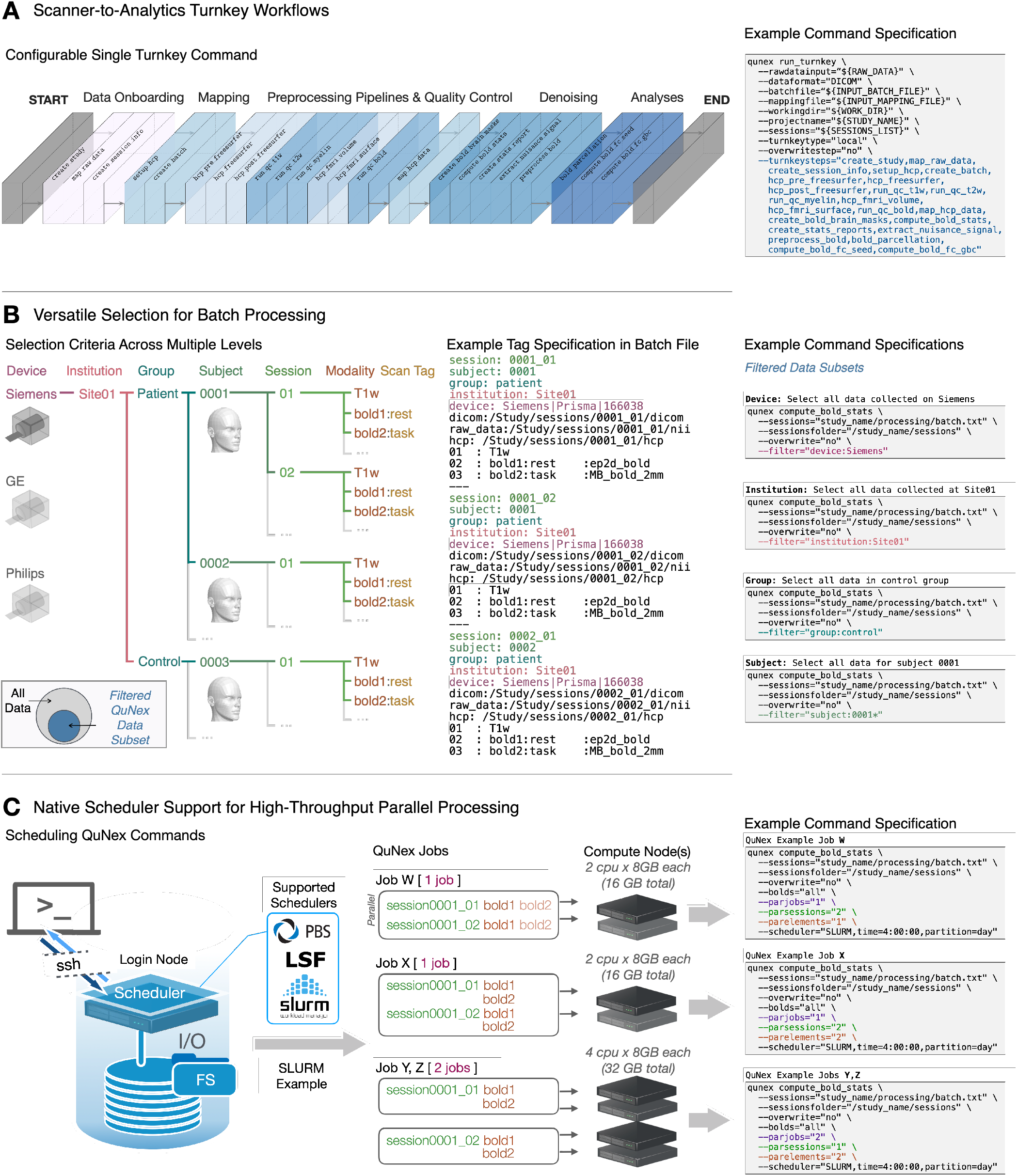
QuNex Turnkey Functionality and Batch Engine for High-throughput Processing. **A)** QuNex provides a “turnkey” engine which enables fully automated deployment of entire pipelines on neuroimaging data via a single command (`qunex run_turnkey`). An example of a typical workflow with key steps supported by the turnkey engine is highlighted, along with the example command specification. QuNex supports state-of-the-art preprocessing tools from the neuroimaging community (e.g. the HCP MPP (9)). For a detailed visual schematic of QuNex steps and commands, see **Supplementary Information** and **Figure S1. B)** The QuNex batch specification is designed to enable flexible and comprehensive “filtering” and selection of specific data subsets to process. The filtering criteria can be specified at multiple levels, such as devices (e.g. Siemens, GE, or Philips MRI scanners), institutions (e.g. scanning sites), groups (e.g. patient vs controls), subjects, sessions (e.g. time points in a longitudinal study), modalities (e.g. T1w, T2w, BOLD, diffusion), or scan tags (e.g. name of scan). **C)** QuNex natively supports job scheduling via LSF, SLURM, or PBS schedulers and can be easily deployed in HPC systems to handle high-throughput, parallel processing of large neuroimaging datasets. The scheduling options enable precise specification of paralellization both across sessions and within session (e.g., parallel processing of BOLD images) for optimal performance and utilization of cluster resources.

### Filtering Grammar Enables Flexible Selection of Study-Specific Data Processing

Flexible selection of sessions/scans for specific steps is an essential feature for dataset management, especially datasets from multiple sites, scanners, participant groups, or scan types. For example, the user may need to execute a command only on data from a specific scanner; or only on resting-state (versus task-based) functional scans for all sessions in the study. QuNex enables such selection with a powerful filtering grammar in the study-level “batch files”, which are text files that are generated as part of the onboarding process.

Batch files contain metadata about the imaging data and various acquisition parameters (e.g., site, device vendor, group, subject ID, session ID, acquired modalities) and serve as a record of all session-specific information in a particular study. When users create the batch file through the `create_batch` command, QuNex sifts through all sessions in the study and adds the information it needs for further processing and analyses to the batch file. This makes the batch file a key hub that stores all the relevant study meta-data. One of the key advantages of this approach is that users can easily execute commands on all or only a specific subset of sessions from a study by filtering the study-level batch file. **Figure 2B** visualizes the logic behind filtering data subsets from batch files and examples of the `filter` parameter in a QuNex command. Information about each scan (e.g., scanner/device, institution/scan site, group, subject ID, session ID, modality, scan tag) in the batch file is provided using a `key:value` format (e.g., `group:patient`). While some keys are required for QuNex processing steps (e.g., `session, subject`) and are populated automatically during the onboarding process, users can add as many additional `key:value` tags as they need. The `filter` parameter in a QuNex command will search through the batch file and select only the scans with the specified `key:value` tag. This filtering can be executed at multiple levels, from selecting all scans from a particular type of scanner to scans from only a single session. For example, the setting `filter=“device:Siemens”` will select all data for scans conducted by a Siemens scanner, whereas the setting `filter=“session:0001_1”` will select only data from the session ID 0001_1.

### QuNex Provides Native Scheduler Support for Job Management

Many institutions use HPC systems or cloud-based servers for processing, necessitating job management applications such as scheduler software and custom scheduling scripts (see examples in **Supplementary Information** and **Figure S5**). This is especially important for efficient processing of large datasets which may include thousands of sessions. While QuNex is platform-agnostic, all QuNex commands, including `run_turnkey`, are compatible with commonly used scheduling systems (i.e., SLURM, PBS and LSF) for job management in HPC systems (**Figure S6**). Thus, QuNex is easily scalable and equipped to handle high-throughput, parallel processing of large neuroimaging datasets. To schedule a command on a cluster, users simply provide a `scheduler` parameter to any QuNex command call and the command will be executed as a job on an HPC system, eliminating the need for specialized scripts with scheduling directives. Additionally, QuNex provides parameters for users to easily customize the parallelization of their jobs from the command line call. The `parjobs` parameter specifies the total number of jobs to run in parallel; `parsessions` specifies the number of sessions to run in parallel within any single job; and `parelements` specifies the number of elements (e.g., fMRI runs) within each session to run in parallel. Users can provide the scheduling specification for their jobs to ensure that computational resources are allocated in a specific way; otherwise, QuNex will automatically assign scheduling values for job parallelization, as described in **Figure S7. Figure 2C** shows examples of how the native support for scheduling and QuNex’s parallelization parameters can be leveraged to customize the way processing is distributed across jobs. For example, specifying `parjobs=1, parsessions=2`, and `parelements=1` will ensure that only one job is run at a time on the compute nodes, with two sessions running in parallel. Any individual elements within each session (e.g., multiple BOLD runs) will run serially, one at a time. This parallelization and scheduling functionality, in combination with the turnkey engine and batch specification, is extremely powerful at handling large-scale datasets, while providing great flexibility and user friendliness in optimization to maximally utilize computing resources. Through a single QuNex command line call, a user can onboard, process, and analyze thousands of scans on an HPC system in a parallel manner, drastically reducing the amount of time and effort for datasets of scale.

### Parameter Specification Environment Enables Reproducible Workflows of Multi-modal Datasets

The diversity of neuroimaging parameters can lead to challenges in replicating preprocessing choices and thus affect the reproducibility of results. QuNex supports consistent specification and documentation of parameter values by storing this information in the parameter header of batch files (see **Figure 3B** for an example). Many parameters in neuroimaging pipelines are the same across different steps or commands, or across different command executions (e.g., if data for the same study/scanner are processed sequentially). By providing these parameters and their values in the batch files, users are assured that shared parameters will use the same value across pipeline steps. Furthermore, such specification enables complete transparency and reproducibility, as processing workflows can be fully replicated by using the same batch files, and the batch files themselves can be easily shared between researchers. For convenience, an alternative way of providing parameters is through the CLI call; if a parameter is defined both in the batch file and in the CLI call, the version in the CLI call takes precedence.

**Fig. 3.**
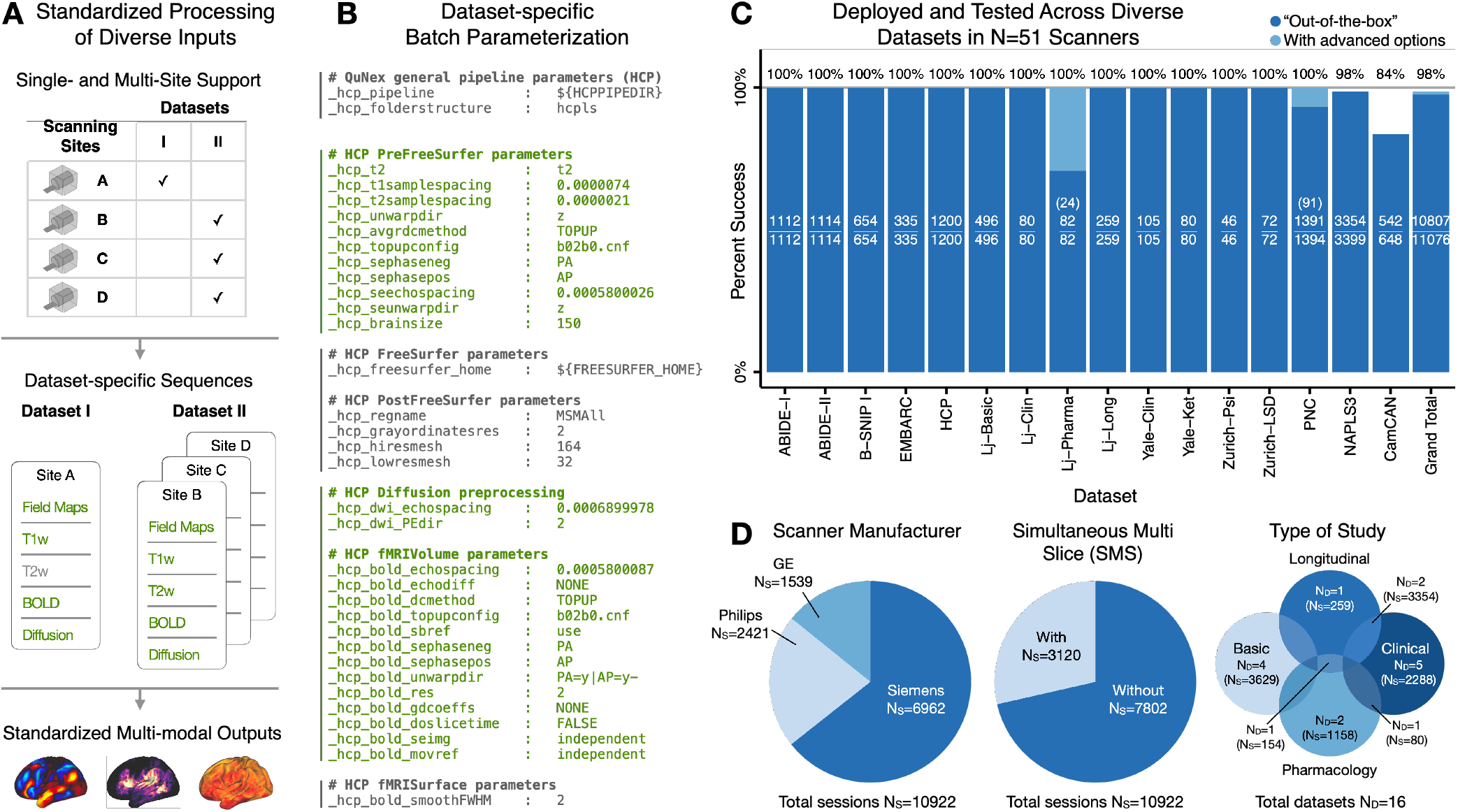
Consistent Processing at Scale and Standardized Outputs Through Batch Specification. **A)** The batch specification mechanism in QuNex is designed to support data processing from single-site and multi-site datasets to produce standardized outputs. Acquisition parameters can be flexibly specified for each sequence. Here, example datasets I (single-site study) and II (multi-site study) illustrate possible use cases, with the sequences in each dataset shown in green text. Although Dataset I does not include T2w scans, and Dataset II contains data from different scanners, all these data can be consistently preprocessed in all modalities to produce standardized output neural features. **B)** Parameters can be tailored for each study in the header of the batch processing file. An example is shown with parameters in green text tailored to Site B in Dataset II (similar to those used in HCP datasets (9)). **C)** QuNex has been highly successful in preprocessing data from numerous publicly available as well as private datasets, totalling over 10,000 independent scan sessions from over 50 different scanners. In some cases, advanced user options (such as custom brain masks) can be used to rescue sessions which failed with “out-of-the-box” default preprocessing options. The number of successful/total sessions is reported in each bar. The number of sessions rescued with advanced options is shown in parentheses, when applicable. The total proportion of successfully preprocessed sessions from each study (including any sessions rerun with advanced options) as well as the grand total across all studies is shown above the bar plots. **D)** QuNex has been successfully used to preprocess data with a wide range of parameters and from diverse datasets. (Left) QuNex has been tested on MRI data acquired with the three major scanner manufacturers (Philips, GE and Siemens). Here *N*_*S*_ specifies the number of individual scan sessions that were acquired with each type of scanner. (Middle) QuNex is capable of processing images acquired both with and without simultaneous multi-slice (SMS) acquisition (also known as multi-band acquisition, i.e.: Simultaneous Multi-Slice in Siemens scanners; Hyperband in GE scanners; and Multi-Band SENSE in Philips scanners (20)). (Right) QuNex has been tested on data from clinical, pharmacology, longitudinal and basic population-based datasets. Here, *N*_*D*_ specifies the number of datasets; *N*_*S*_ specifies the total number of individual scan sessions in those datasets.

Preprocessing functions are typically executed on multiple sessions at the same time so that they can run in parallel. As mentioned above, QuNex utilizes batch files to define processing parameters, in order to facilitate batch processing of sessions. This batch file specification allows QuNex to produce standardized outputs from data across different studies while allowing for differences in acquisition parameters (e.g., in a multi-site study, where scanner manufacturers may differ across sites). **Figure 3A** illustrates two example use-case datasets (Datasets I and II). The flexibility of the QuNex batch parameter specification enables all data from these different studies and scanners to be preprocessed consistently and produce consistent outputs in all modalities. **Figure 3B** illustrates an example of a real-world batch parameter specification. This information is included in the header of a batch file, and is followed by the session-level information (as shown in **Figure 2B**) for all sessions.

We have successfully used QuNex to preprocess and analyze data from a large number of public and private neuroimaging datasets (**Figure 3C**) (4), totalling more than 10,000 independent scan sessions from over 50 different scanners. **Figure 3D** shows that the data differ in terms of the scanner manufacturer (Philips, GE or Siemens), acquisition technique (simultaneous multi-slice/multi-band), and the study purpose (clinical, basic, longitudinal and pharmacology studies). These datasets also span participants from different stages of development, from children to older adults. Across these diverse datasets, the percentage of successfully processed sessions is extremely high: 100% in the majority of studies and ∼98.5% in total across all studies (**Figure 3C**). Of note, QuNex supports the preprocessing efforts of major neuroimaging consortia and is used by the Connectome Coordination Facility (CCF) to preprocess all Lifespan and Connectomes Related to Human Disease (CRHD) datasets (4).

### QuNex Supports Extraction of Multi-modal Features at Multiple Spatial Scales

Feature engineering is a critical choice in neuroimaging studies and features can be computed across multiple spatial scales. Importantly, given the challenges with mapping reproducible brain-behavioral relationships (3), selecting the right features at the appropriate scale is vital for optimizing signal-to-noise in neural data and producing reproducible results. QuNex enables feature generation and extraction at different levels of resolution (including “dense” full-resolution, parcels, or whole-brain networks) for both volume and CIFTI (combined surface and volume) representations of data, consistently across multiple modalities, for converging multi-modal neuroimaging analytics. While some parcellations are currently distributed with QuNex (such as the HCP-MMP1.0 (23), CAB-NP (22) and atlases distributed within FSL/FreeSurfer) users are free to provide and use their own parcellation. **Figure 4** shows convergent multi-modal results in a sample of N=339 unrelated young adults. Myelin (T1w/T2w) maps reflect high myelination in sensorimotor areas such as primary visual and sensorimotor networks, and lower myelination in higher-order association networks (**Figure 4A**) (24). DMRI measures capture the white matter connectivity structure through tract termination (14) and maximal intensity projection (MIP) of the left arcuate fasciculus (**Figure 4B**); as well as structural connectivity (23). For example, seed-based structural connectivity of Broca’s area (26, 27) highlights connections to canonical language areas such as Wernicke’s area (28), superior temporal gyrus and sulcus (29, 30), and frontal language regions (27, 31) (**Figure 4C**). This is consistent with the results of seed-based functional connectivity of Broca’s area from resting-state fMRI data in the same individuals (**Figure 4D**); and furthermore, it is aligned with the activation patterns from a language task (**Figure 4E**) (25). Across modalities, QuNex supports the extraction of metrics as raw values (e.g., Pearson’s r or Fisher’s Z for functional connectivity; probabilistic tractography streamline counts for structural connectivity; t-values for task activation contrasts) or standardized Z-scores.

Notably, features across all modalities can be extracted in a consistent, standardized format after preprocessing and post-processing within QuNex. This enables frictionless comparison of features across modalities, e.g. for multi-modal, multivariate analyses.

### QuNex Enables Single-Session Modeling of Time-series Modalities

Modeling of time-series data, such as BOLD, at the single-session level can be used for a variety of purposes, including nuisance regression and extracting task activation for individual subjects. QuNex supports denoising and modeling of time-series data at the single-session level via a general linear model (GLM) framework, executed through the `preprocess_conc` command. Here, we demonstrate this framework with functional BOLD time-series. **Figure 5A** showcases a use case where resting-state BOLD data are first denoised and then used to compute seed-based functional connectivity maps of the primary somatosensory area (S1). During the denoising step, the user can choose which sources of nuisance signal to remove (including motion parameters and their derivatives and BOLD signals extracted from ventricles, white matter, whole brain or any other custom defined regions, and their first derivatives). These nuisance signals are included as covariates in the GLM, which produces, for each BOLD run, residual time-series data as well as coefficient maps for all specified regressors. The denoised time-series can then be used for further analytics, e.g. by computing seed-based functional connectivity using the `fc_compute_seedmaps` command.

**Fig. 4.**
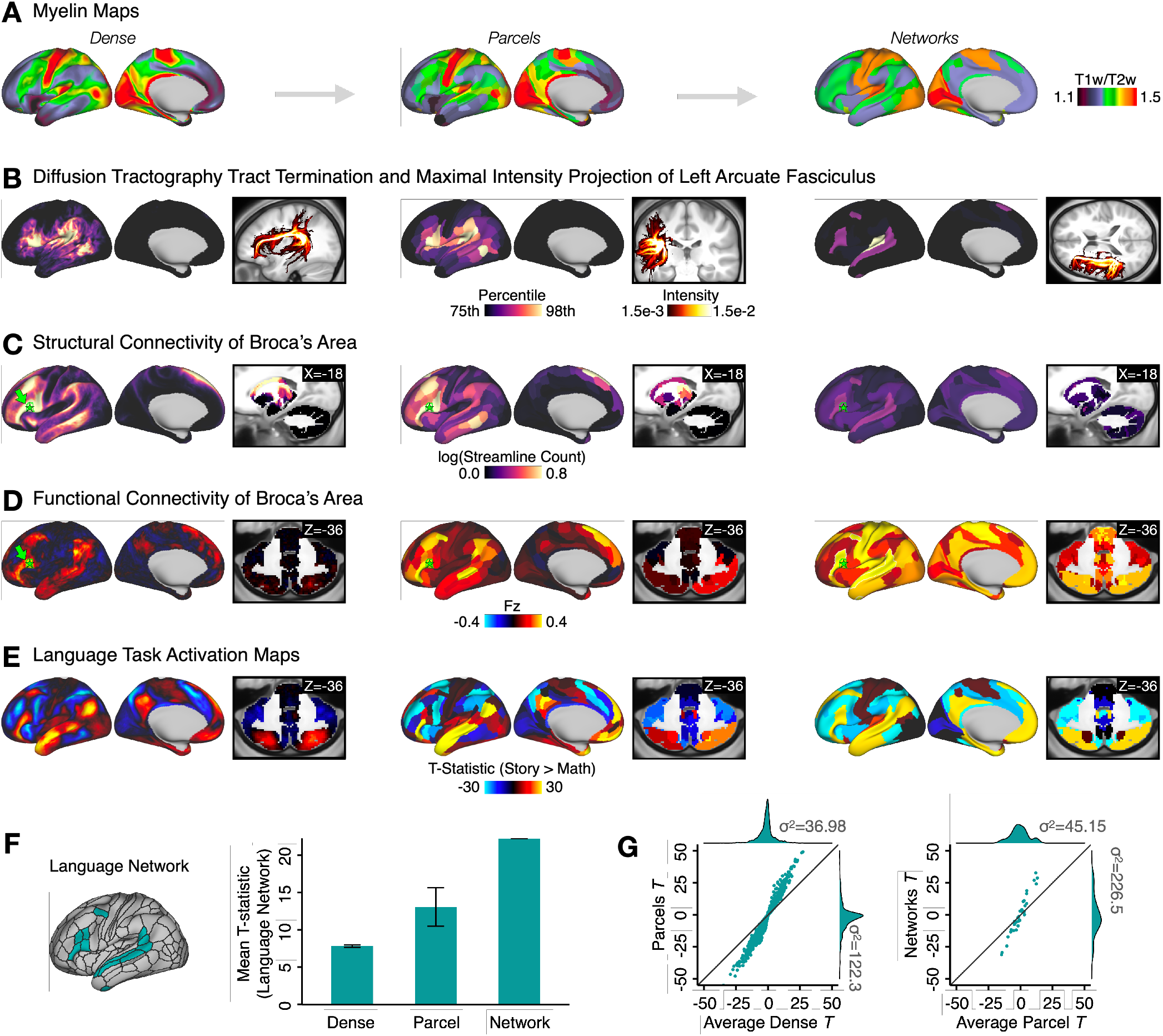
Extracting Multi-modal Processing Features at Multiple Levels of Resolution. Output features from multiple modalities are shown, as an example of a crossmodal analysis that may be done for a study. Here, features were computed from a cohort of N=339 unrelated subjects from the HCP Young Adult cohort (21). In addition to cross-modality support, QuNex offers feature extraction at “dense” (i.e. full-resolution), parcel-level and network-level resolutions. All features are shown below at all three resolutions. We used the Cole-Anticevic Brainwide Network Parcellation (CAB-NP) (22, 23), computed using resting-state functional connectivity from the same cohort and validated and characterized extensively in (22). **A)** Myelin maps, estimated using the ratio of T1w/T2w images (24). **B)** Left arcuate fasciculus computed via diffusion tractography (14). Surface views show the cortical tract termination (white-grey matter boundary endpoints) and volume views show the maximal intensity projection. **C)** Structural connectivity of Broca’s area (parcel corresponding to Brodmann’s Area [BA] 44, green star) (23). **D)** Resting-state functional connectivity of Broca’s area (green star). For parcel- and network-level maps, resting-state data were first parcellated before computing connectivity. **E)** Task activation maps for for the “Story versus Math” contrast in a language processing task (25). For parcel- and network-level maps, task fMRI data were first parcellated before model fitting. **F)** Left: Whole-brain Language network from the CAB-NP (22). (Right) The mean t-statistic within Language network regions from the “Story versus Math” contrast (shown in panel E) improves when data are first parcellated at the parcel-level relative to dense-level data and shows the greatest improvement when data are first parcellated at the network-level. Error bars show the standard error. **G)** (Left) T-statistics computed on the average parcel beta estimates are higher compared to the average T-statistics computed over dense estimates of the same parcel. Teal dots represent 718 parcels from the CAB-NP × 3 Language task contrasts (“Story versus Baseline’’; “Math versus Baseline’’; “Story versus Math”). (Right) Similarly, T-statistics computed on beta estimates for the network are higher than the average of T-statistics computed across parcels within each network.

**Fig. 5.**
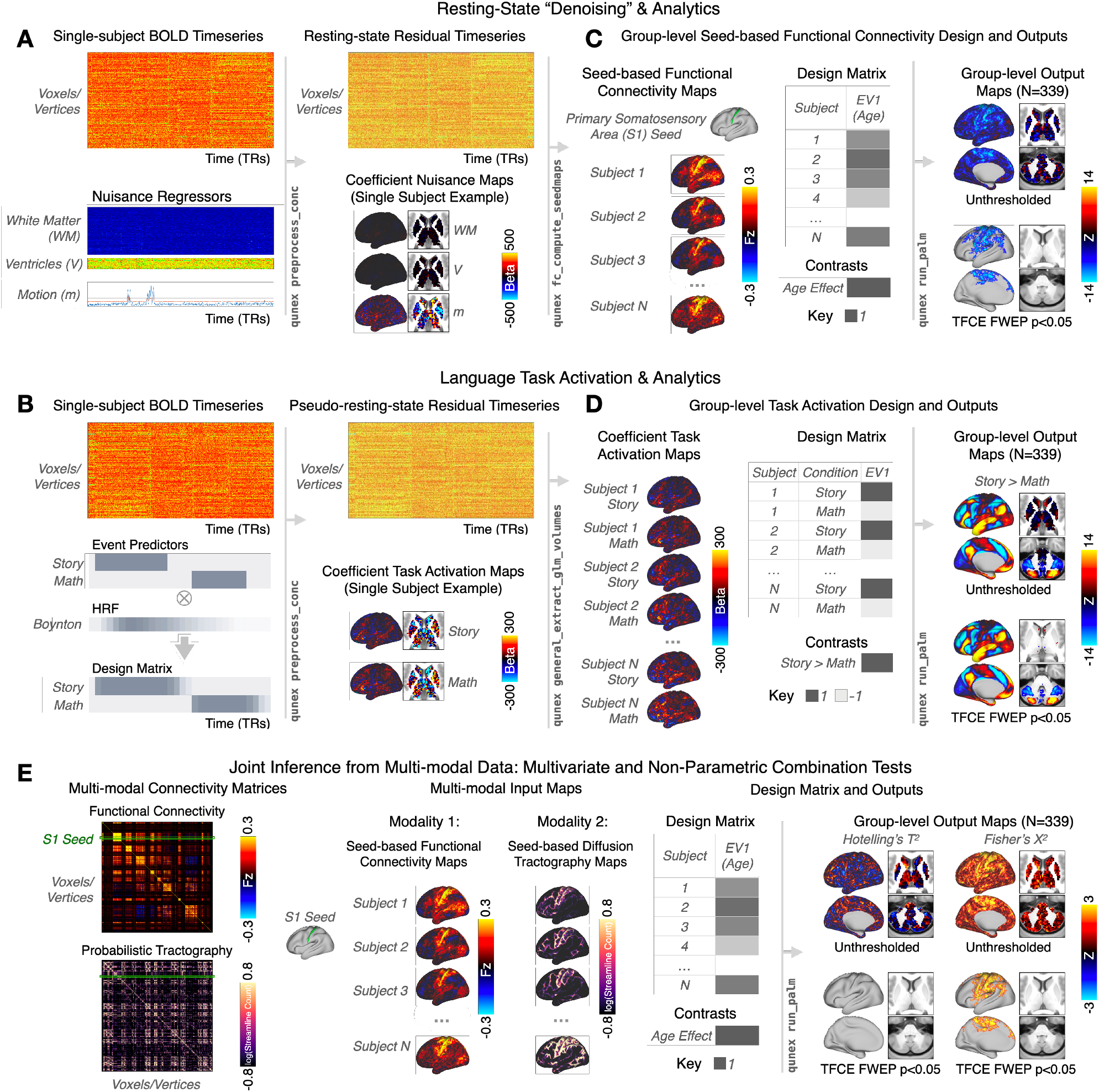
General Linear Model (GLM) for Single-Session Modeling of Time-series Modalities and Integrated Interoperability with PALM for Group-Level Analytics. **A)** The QuNex GLM framework enables denoising and/or event modeling of resting-state and task BOLD images at the individual-session level in a single step. A use case is shown for resting-state BOLD data. At the single-subject level, individual nuisance regressors (such as white matter and ventricular signal and motion parameters) can be specified such that they are regressed out of the BOLD timeseries with the `qunex preprocess_conc` function. The regressors can be per-frame (as shown), per-trial, or even per-block. The GLM outputs a residual timeseries of “denoised” resting-state data as well as one coefficient map per nuisance regressor. The resting-state data for each subject can then be used to calculate subject-specific feature maps, such as seed-based functional connectivity maps with `qunex fc_compute_seedmaps`. **B)** The GLM engine can also be used for complex modeling and analysis of task events, following a similar framework. Event modeling is specified in `qunex preprocess_conc` by providing the associated event file; the method of modeling can be either assumed (using a hemodynamic response function [HRF]) or unassumed. Here, an example from the HCP’s Language task is shown. The two events, “Story” and “Math”, are convolved with the Boynton HRF to build the subject-level GLM. As with the resting-state use case shown in **A**, the GLM outputs the single-subject residual timeseries (in this case ‘pseudo-resting state’) as well as the coefficient maps for each regressor, here the Story and Math tasks. **C)** Connectivity maps from all subjects can then be entered into a group-level GLM analysis. In this example, the linear relationship between connectivity from the primary somatosensory area (S1) seed and age across subjects is tested in a simple GLM design with one group and one explanatory variable (EV) covariate, demeaned age. QuNex supports flexible group-level GLM analyses with non-parametric tests via Permutation Analysis of Linear Models (PALM, (12)), through the `qunex run_palm` function. The specification of the GLM and individual contrasts is completely configurable and allows for flexible and specific hypothesis testing. Group-level outputs include full uncorrected statistical maps for each specified contrast as well as p-value maps that can be used for thresholding. Significance for group-level statistical maps can be assessed with the native PALM support for TFCE ((12), shown) or cluster statistics with familywise error protection (FWEP). **D)** The subject-level task coefficient maps can then be input into the `qunex run_palm` command along with the group-level design matrix and contrasts. The group-level output maps show the differences in activation between the Story and Math conditions. **E)** QuNex also supports multi-variate and joint inference tests for testing hypotheses using data from multiple modalities, such as BOLD signal and dMRI. Example connectivity matrices are shown for these two modalities, with the S1 seed highlighted. Similar to the use cases shown above, maps from all subjects can be entered into a group-level analysis with a group-level design matrix and contrasts using the `qunex run_palm` command. In this example, the relationship between age and S1-seeded functional connectivity and structural connectivity is assessed using a Hotelling’s *T*^2^ test and Fisher’s *X*^2^. The resulting output maps show the unthresholded and thresholded (p < 0.05 FWEP, 10,000 permutations) relationship between age and both neural modalities.

For task data, QuNex facilitates the building of design matrices at the single session-level (**Figure 5B**). The design matrices can include: i) task regressors created by convolving a haemodynamic response function (HRF, e.g. Boynton, double Gaussian) with event timeseries (i.e. assumed modeling, as shown here for the Story and Math blocks of a language task (25)); ii) separate regressors for each frame of the trial, i.e. unassumed modeling of task response; iii) a combination of assumed and unassumed regressors. The events in assumed and unassumed modeling can be individually weighted, enabling estimates of trial-by-trial correlation with e.g. response reaction time, accuracy or precision. The GLM engine estimates the model and outputs both a residual time-series (“pseudo-resting state”) as well as coefficient maps for each regressor, reflecting task activation for each of the modelled events. After a model has been estimated, it is possible to compute both predicted and residual timeseries with an arbitrary combination of regressors from the estimated model (e.g., residual that retains transient task response after removal of sustained task response and nuisance regressors).

### QuNex Supports Built-In Interoperability with Externally-Developed Tools

QuNex is designed to provide interoperability between community tools to remove barriers between different stages of neuroimaging research. One such feature is its compatibility with XNAT (eXtensible Neuroimaging Archive Toolkit) (32, 33), a widely used platform for research data transfer, archiving, and sharing (**Figure S8**). This enables reseachers to seamlessly organize, process, and manage their imaging studies in a coherent integrated environment. QuNex also provides user-friendly interoperability with a suite of tools, including AFNI, FSL, HCP Workbench etc.

Another interoperabilty feature is the execution of group-level statistical testing of neuroimaging maps, which is performed through Permutation Analysis of Linear Models (PALM) (12), an externally-developed tool which executes nonparametric permutation-based significance testing for neuroimaging data. QuNex provides a smooth interface for multi-level modeling via PALM. PALM itself supports volume-based NIFTI, surface-based GIFTI, and surface-volume hybrid CIFTI images, and allows for fully customizable statistical tests with a host of familywise error protection and spatial statistics options. Within QuNex, PALM is called through the `qunex run_palm` command, which provides a cohesive interface for specifying inputs, outputs, and options. The user is able to customize design matrices and contrasts according to their need and provide these along with QuNex-generated neural maps to assess for significance using permutation testing and familywise error protection.

**Figure 5C** illustrates an example where S1-seed functional connectivity maps for N=339 sessions are tested at the group-level to show a significant negative relationship with age in areas such as the somatomotor cortices (p<0.05, nonparametrically tested and family-wise error protected with threshold-free cluster enhancement (TFCE) (34)). As with functional connectivity maps, task activation maps can be tested for significant effects in the group-level GLM with PALM **Figure 5D**. Here, a within-subject t-test of the Story > Math contrast reveals significant areas of the language network, also shown in **Figure 4E-F**. QuNex additionally supports joint inference from combined multi-modal data via multivariate statistical tests (e.g., MANOVAs, MANCOVAs) and non-parametric combination tests (35), also executed through PALM and thus compatible with permutation testing. For example, seed-based functional connectivity and structural connectivity of area S1 from the same individuals can be entered into the same test as separate modalities. The second-level GLM shown in **Figure 5E** is the same one as in **Figure 5B** to test for age effects. Such joint inference tests can be used to test whether there are jointly significant differences on a set of modalities. Thus, QuNex enables streamlined workflows for multi-modal neuroimaging feature generation and integrated multi-variate statistical analyses. QuNex workflows simplify neuroimaging data management and analysis across a wide range of clinical, translational, and basic neuroimaging studies, including studies examining the relationship between neuroimaging features and gene expression or symptom presentation, or pharmacological neuroimaging studies of mechanism. **Figure S9** highlights a few examples of recently published studies which leveraged QuNex for preprocessing, feature generation, and analytics.

QuNex also encourages future integration of open source community tools via the extensions framework, through which researchers can integrate their own tools and pipelines into the QuNex platform (**Supplementary Information**). To continually engage community participation in neuroimaging tool development, QuNex provides a SDK that includes helper functions for users to set up a development and testing environment (**Figure S10**).

### Cross-Species Support for Translational Neuroimaging

Studies of non-human species have substantially contributed to the understanding of the central nervous system, and provided a crucial opportunity for translational science. In particular, the macaque brain is phylogenetically similar to the human brain, and comparative neuroimaging studies in macaques have served to inform and validate human neuroimaging results. It is thus imperative to develop and distribute tools for consistent processing and analytics of non-human neuroimaging data for aiding translational cross-species neuroimaging studies (36, 37). To this end, QuNex supports analogous workflows for human and non-human primate neuroimaging data. **Figure 6** shows parallel steps for running HCP-style preprocessing and generating multimodal neural features in human and macaque data. Structural data outputs include FreeSurfer segmentation and labelling of cortical and subcortical areas, T1w/T2w myelin maps (**Figure 6A**), and structural metrics such as cortical thickness, curvature, and subcortical volumes. Functional data outputs include BOLD signal and metrics such as functional connectivity (**Figure 6B**). Diffusion metrics include measures of microstructure (e.g. fractional anisotropy maps), white matter tracts and their cortical termination maps, and whole-brain structural connectivity, as shown in **Figure 6C**. Currently, QuNex supports macaque diffusion pipelines in the released container, with HCP macaque functional neuroimaging pipelines (18) and mouse neuroimaging pipelines (19) under development for a future release. The functional macaque images shown here are obtained from an early development version of the pipelines.

**Fig. 6.**
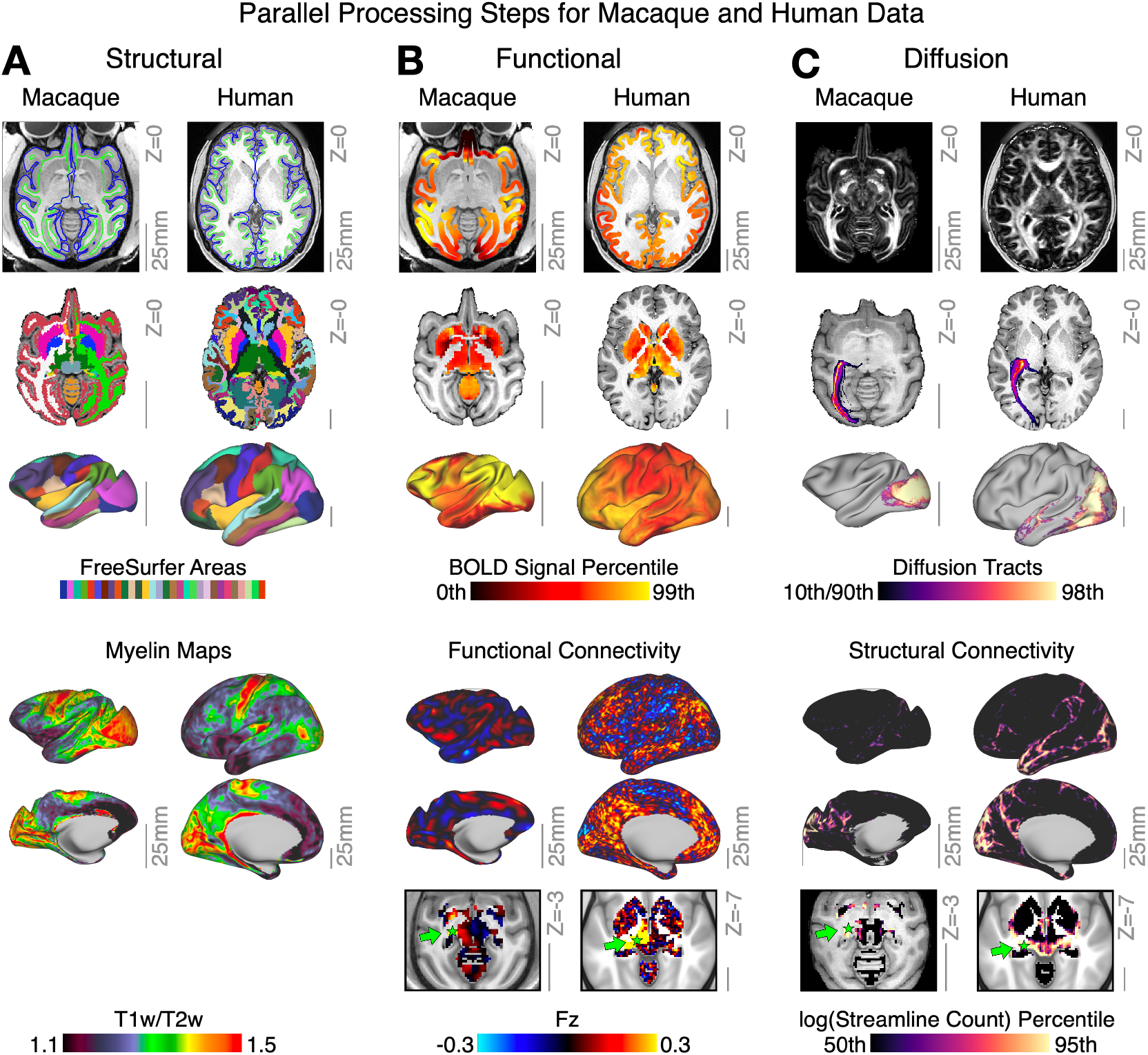
QuNex Enables Neuroimaging Workflows Across Different Species. **A)** Structural features for exemplar macaque and human data, including surface reconstructions and segmentation from FreeSurfer. Lower panel shows output myelin (T1w/T2w) maps. **B)** Functional features for exemplar macaque and human showing BOLD signal mapped to both volume and surface. Lower panels show and resting-state functional connectivity seeded from the lateral geniculate nucleus of the thalamus (green arrow). **C)** Diffusion features for exemplar macaque and human data, showing whole-brain fractional anistropy, and volume and surface terminations of the left optic radiation tract. Lower panels show the structural connectivity maps seeded from the lateral geniculate nucleus of the thalamus (green arrow). Grey scale reference bars in each panel are scaled to 25mm.

## Discussion

The popularity of neuroimaging research has led to the development and availability of many tools and pipelines, many of which are specific to one modality. This in turn has led to challenges in method integration, particularly across different neuroimaging sub-fields. Additionally, the wide availability of different pipeline and preprocessing/analytic choices may contribute to difficulties with producing replicable results (13). Thus, QuNex is designed to be an integrative platform with interoperability for externally-developed tools across multiple neuroimaging modalities. It leverages existing state-of-the-art neuroimaging tools and software packages, with a roadmap for continued integration of new tools and features. Additionally, QuNex provides features such as turnkey functionality, native scheduler support, flexible data filtering and selection, multi-modal integration, and crossspecies support, to fully enable neuroimaging workflows.

It should be noted that there are currently several tools in the neuroimaging community with multi-modality support, including (but not limited to) FSL, SPM, Freesurfer, AFNI, and PALM. These softwares all offer preprocessing and/or analytic capalities for at least 3 different neural modalities, such as T1w, T2w, fMRI, arterial spin labelling (ASL), dMRI, EEG, MEG, and functional near-infrared spectroscopy (fNIRS). Rather than reinventing the wheel, QuNex builds upon the decades of research, optimization, and validation of these tools by using them as basic building blocks for fundamental steps of neuroimaging workflows, and augments their functionality and interoperability. Other highlevel environments, such as HCP MPP (9), UK Biobank pipelines (38), fMRIPrep (10), QSIPrep (11), micapipe (39), nipype (17), BrainVoyager (40), FuNP (41), Clinica (42), brainlife (43), NeuroDebian (44), and LONI pipelines (45), also leverage other neuroimaging tools as building blocks. Many of these options are uni-modal preprocessing pipelines (e.g., fMRIPrep, QSIPrep) or preprocessing pipelines developed for specific consortia (HCP and UKBiobank pipelines). We emphasize that QuNex is a unifying framework for integrating multi-modal, multi-species neuroimaging tools and workflows, rather than a choice of preprocessing or analytic pipeline; as such, QuNex can incorporate these options, as evidenced by the current integration of the HCP MPP and the planned integration of fMRIPrep. Furthermore, QuNex offers additional user-friendly features which expand upon the existing functionality of these tools, including flexible data filtering, turnkey functionality, support for cloud and HPC deployment, native scheduling and parallelization options, and collaborative development tools. A list of the implementations for different functionalities in QuNex, as well as comparable implementations in other neuroimaging pipelines and environments, is shown in **Figure S11**.

In addition, several commercial platforms are available for neuroimaging data management and analytics (e.g., Flywheel (46), QMENTA, Nordic Tools, Ceretype), especially for clinical applications. While these platforms offer a wide range of neuroinformatics functionalities, they are difficult to evaluate due to their high cost of services and proprietary content. On the contrary, QuNex is free to use for non-commercial research, with transparent and collaborative code and development.

The QuNex container and SDK are available at qunex.yale.edu. Interested parties can register for access and download QuNex as well as as example data and tutorials. The online documentation can be found at http://qunex.readthedocs.io/ and the community forum is hosted at forum.qunex.yale.edu.

Neuroimaging is an actively advancing field and QuNex is committed to continual development and advancement of neuroimaging methods. Below, we list features and existing external software which are currently under development/integration, as well as those which are staged for future release. As neuroimaging techniques advance and novel tools and methods are developed and adopted, we plan to integrate them into the QuNex platform either through internal development or via the extensions framework.

### Currently under development

Longitudinal preprocessing; mouse neuroimaging preprocessing and analytics; EEG preprocessing and analytics.

### Staged for development

PET preprocessing and analytics; BIDS exporter; fMRIPrep.

## Acknowledgments

We would like to thank Brendan Adkinson, Charles Schleifer, Martina Starc, Anka Slana Ozimič, and Martin Bavčar for their support in testing the QuNex platform, and contributing to the development of QuNex documentation. Financial support for this study was provided by NIH grants DP5OD012109-01 (to A.A.), 1U01MH121766 (to A.A.), R01MH112746 (to J.D.M.), 5R01MH112189 (to A.A.), 5R01MH108590 (to A.A.), NIAAA grant 2P50AA012870-11 (to A.A.), NSF NeuroNex grant 2015276 (to J.D.M.), the Brain and Behavior Research Foundation Young Investigator Award (to A.A.), SFARI Pilot Award (to J.D.M. & A.A.), BlackThorn Therapeutics (to J.D.M. & A.A.), the European Research Council (Consolidator Grant 101000969 to S.N.S. and S.W.), Wellcome Trust (Grant 217266/Z/19/Z to S.N.S.), the Slovenian Research Agency (ARRS) (Grant Nos. J7-8275, J7-6829, P3-0338 to G.R.).

## Author Contributions

J.L.J., J.D., S.W., S.N.S., A.A., and G.R. prepared the initial blockout of the manuscript. J.L.J. and J.D. prepared the figures and the initial draft of the manuscript. A.A. and G.R. supervised this research. All authors helped with contributed to the development of the QuNex platform. All authors reviewed and approved the final version of the manuscript.

## Methods

### Description of the Preprocessing Validation Datasets

We tested preprocessing using QuNex on a total of 16 datasets, including both publicly-available and aggregated internal datasets. For each dataset, we prepared batch files with parameters specific to the study (or site, if the study is multisite and acquisition parameters differed between sites). We then used QuNex commands to run all sessions through the HCP Minimal Preprocessing Pipelines (MPP) for structural (T1w images; T2w if available), functional data, and diffusion data (if available). A brief description of each dataset in **Figure 3** is in the **Supplementary Information**. Additional details on diffusion datasets and preprocessing can also be found below.

### Preprocessing of Validation Datasets

All datasets were preprocessed using QuNex with the HCP MPP (9) (`qunex hcp_pre_freesurfer; hcp_freesurfer; hcp_post_freesurfer; hcp_fmri_volume; hcp_fmri_surface`). A summary of the HCP Pipelines is as follows: the T1w structural images were first aligned by warping them to the standard Montreal Neurological Institute-152 (MNI-152) brain template in a single step, through a combination of linear and non-linear transformations via the FMRIB Software Library (FSL) linear image registration tool (FLIRT) and non-linear image registration tool (FNIRT) (47). If a T2w was present, it was co-registered to the T1w image. If field maps were collected, these were used to perform distortion correction. Next, FreeSurfer’s recon-all pipeline was used to segment brain-wide gray and white matter to produce individual cortical and subcortical anatomical segmentations (48). Cortical surface models were generated for pial and white matter boundaries as well as segmentation masks for each subcortical grey matter voxel. The T2w image was used to refine the surface tracing. Using the pial and white matter surface boundaries, a ‘cortical ribbon’ was defined along with corresponding subcortical voxels, which were combined to generate the neural file in the Connectivity Informatics Technology Initiative (CIFTI) volume/surface ‘grayordinate’ space for each individual subject (9). BOLD data were motion-corrected by aligning to the middle frame of every run via FLIRT in the initial NIFTI volume space. Next a brain-mask was applied to exclude signal from non-brain tissue. Next, cortical BOLD data were converted to the CIFTI gray matter matrix by sampling from the anatomically-defined gray matter cortical ribbon and subsequently aligned to the HCP atlas using surface-based nonlinear deformation (9). Subcortical voxels were aligned to the MNI-152 atlas using whole-brain non-linear registration and then the Freesurfer-defined subcortical segmentation was applied to isolate the CIFTI subcortex. For datasets without field maps and/or a T2w image, we used a version of the MPP adapted for compatibility with “legacy” data, featured as a standard option in the HCP Pipelines provided by the QuNex team (https://github.com/Washington-University/HCPpipelines/pull/156). The adaptations for single-band BOLD acquisition have been described in prior publications (49, 50). Briefly, adjustments include allowing the HCP MPP to be conducted without high-resolution registration using T2w images and without optional distortion correction using field maps. For validation of preprocessing via QuNex, we counted the number of sessions in each study which successfully completed the HCP MPP versus the number of sessions which errored during the pipeline.

### Description of the Datasets Used for Analytics

#### HCP Young Adults (HCP-YA) Dataset

To demonstrate neuroimaging analytics and feature generation in human data, we used N=339 unrelated subjects from the HCP-YA cohort (21). The functional data from these subjects underwent additional processing and removal of artifactual signal after the HCP MPP. These steps included ICA-FIX (9, 51) and movement scrubbing (52) as done in our prior work (50, 53). We combined the four 15-min resting-state BOLD runs in order of acquisition, after first demeaning each run individually and removing the first 100 frames to remove potential magnetization effects (22). Seed-based functional connectivity was computed using `qunex fc_compute_seedmaps` and calculated as the Fisher’s Z-transformed Pearson’s r-value between the seed region BOLD time-series and time-series in the rest of brain. Task activation maps were computed from a language processing task (25), derived from (54). Briefly, the task consisted of two runs, each with 4 blocks of 3 conditions: (i) Sentence presentation with detection of semantic, syntactic and pragmatic violations; (ii) Story presentation with comprehension questions (‘Story’ condition); (iii) Math problems involving sets of arithmetic problems and response periods (‘Math’ condition). Trials were presented auditorily and participants chose one of two answers by pushing a button. Task-evoked signal for the Language task was computed by fitting a GLM to preprocessed BOLD time series data with `qunex preprocess_conc`. Two predictors were included in the model for the ‘Story’ and ‘Math’ blocks, respectively. Each block was approximately 30s in length and the sustained activity across each block was modeled using the Boynton HRF (55). Results shown here are from the Story versus Math contrast (22, 23). Across all tests, statistical significance was assessed with PALM (12) via `qunex run_palm`. Briefly, threshold-free cluster enhancement was applied (34) and the data were randomly permuted 5,000 times to obtain a null distribution. All contrasts were corrected for family-wise error. Diffusion data from this dataset were first preprocessed with the HCP MPP (9) via `qunex hcp_diffusion`, including susceptibility and eddy-current induced distortion and motion correction (56, 57) and the estimation of dMRI to MNI-152 (via the T1wandersson2016integrated space) registration fields. Next, fiber orientations were modelled for up to three orientations per voxel using the FSL’s bedpostX crossing fibers diffusion model. (58, 59), via `qunex dwi_bedpostx_gpu`. After registering to the standard space, whole brain probabilistic tractography was run with FSL’s probtrackx via `qunex dwi_probtracx_dense_gpu`, Producing a dense connectivity matrix for the full CIFTI space. Further, we estimated 42 white matter fibre bundles, and their cortical termination maps, for each subject via XTRACT (14). Following individual tracking, resultant tracts were group-averaged by binarizing normalized streamline path distributions at a threshold and averaging binary masks across the cohort to give the percentage of subjects for which a given tract is present at a given voxel. For all tracts except the middle cerebellar peduncle (MCP), which is not represented in CIFTI surface file formats, the cortical termination map was estimated using connectivity blueprints, as described in (60). These maps reflect the the termination points of the corresponding tract on the white-grey matter boundary surface.

#### Non-human Primate Macaque Datasets

Neural data from two macaques (one in vivo, one ex vivo) are shown. Structural (T1w, T2w, myelin) and functional BOLD data were obtained from a session in the publicly-available PRIMatE Data Exchange (PRIME-DE) repository (61), specifically from the University of California-Davis dataset. In this protocol, subjects were anesthesized with ketamine, dexmedetomidine, or buprenorphine prior to intubation and placement in stereotaxic frame with 1-2% isoflurane maintenance anesthesia during the scanning protocol. They underwent 13.5 min of resting-state BOLD acquisition (gradient echo voxel size: 1.4×1.4×1.4mm; TE: 24ms; TR: 1600ms; FOV = 140mm) as well as T1w (voxel size: 0.3×0.3×0.3mm; TE: 3.65ms; TR: 2500ms; TI: 1100ms; flip angle: 7°), T2w (voxel size: 0.3×0.3×0.3mm; TE: 307ms; TR: 3000ms), spin-echo field maps, and diffusion on a Siemens Skyra 3T scanner with a 4-channel clamshell coil. Preprocessing steps are consistent with the HCP MPP and described in detail in (18, 62).

The high-resolution macaque diffusion data shown were obtained ex vivo and have been previously described (14, 60, 63) and are available via PRIME-DE (http://fcon_1000.projects.nitrc.org/indi/PRIME/oxford2.html). The brains were soaked in phosphate-buffered saline before scanning and placed in fomblin or fluorinert during the scan. Data were acquired at the University of Oxford on a 7T magnet with an Agilent DirectDrive console (Agilent Technologies, Santa Clara, CA, USA) using a 2D diffusion-weighted spin-echo protocol with single line readout (DW-SEMS, TE/TR: 25ms/10s; matrix size: 128×128; resolution: 0.6×0.6mm; number of slices: 128; slice thickness: 0.6mm). Diffusion data were acquired over the course of 53 hours. For each subject, 16 non-diffusion-weighted (b=0s/mm^2^) and 128 diffusion-weighted (b=4000s/mm^2^) volumes were acquired with diffusion directions distributed over the whole sphere. FA maps were reigstered to the standard F99 space (64) using FNIRT. As with the human data, the macaque diffusion data were modelled using the crossing fibre model from bedpostX and used to inform tractography. Again, 42 white matter fibre bundles, and their cortical termination maps, were estimated using XTRACT.

### Functional Parcellation and Seed Definitions

We used the Cole-Anticevic Brain-wide Network Partition (CAB-NP) (22), based on the HCP MMP (23), for definitions of functional networks (e.g. the Language network) and parcels in the cortex and subcortex. Broca’s Area was defined as Brodmann’s Area 44, corresponding to the parcel labelled “L_44_ROI” in the HCP MMP and “Language-14_L-Ctx” in the CAB-NP (23). The left Primary Somatorysensory Area (S1) region was defined as Brodmann’s Area 1 and corresponds to the parcel labelled “L_1_ROI” in the HCP MMP and “Somatomotor-29_L-Ctx” in the CAB-NP (23).

### Design and Features for Open Science

QuNex is developed in accordance to modern standards in software engineering. Adhering to these standards results in a consistently structured, well documented and strictly versioned platform. All QuNex code is open and well commented which both eases and encourages community development. Furthermore, our Git repositories use the GitFlow branching model which, besides keeping our repositories neat and tidy, also helps with the process of merging community developed features into our solution. QuNex has an extensive documentation, both in the form of inline help, accessible from CLI and a Wiki page. Inline documentation offers a short description of all QuNex commands and their parameters while the Wiki documentation offers a number of tutorials and more extensive usage guides. Furthermore, users can establish a direct communication with QuNex developers through the official QuNex forum (https://forum.qunex.yale.edu/), where they can get additional support and discuss or suggest possible new features or anything else QuNex related. To assure maximum possible levels of tractability and reproducibility, QuNex is versioned by using the semantic versioning process (https://semver.org/). The QuNex platform is completely free and open source – QuNex source code is licensed under the GPL (GNU General Public License). Furthermore, QuNex is not only open by nature, but also by design. In other words, we did not simply open up the QuNex code base, we developed it to be as open and accessible as possible. To open up QuNex to the neuroinformatics community, we designed a specialized extensions framework. This framework supports development in multiple programming languages (e.g. Python, MATLAB, R, Bash) and was built with the sole intention to ease the integration of custom community based processing and analysis commands into the QuNex platform. Extensions developed through this extensions framework can access all the tools and utilities (e.g. the batch turnkey engine, logging, scheduling …) residing in the core QuNex code. Once developed, QuNex Extensions are seamlessly attached to the QuNex platform and ran in the same fashion as all existing QuNex commands. Our end goal is to fold the best extensions into our core codebase and thus have a community supported, organically growing neuroimaging platform. As mentioned, to ease this process we have also prepared an SDK, which includes the guidelines and tools that should both speed up the extension development process and make extensions code more consistent with the core QuNex code. This will then allow for faster adoption of QuNex Extensions into the core codebase. See **Figure S10** for visualization of the QuNex Extensions framework.

Since QuNex and other similar platforms depend on a number of software tools which are developed independently, assuring complete reproducibility can be a challenging task since researchers are required to track and archive all the dependencies. To alleviate this issue we publish a container along each unique QuNex version. As a result, using the container for processing and analysis allows users to achieve complete reproducibility by tracking a single number – the version of the QuNex platform used in processing and analysis. QuNex containers are not only important because they offer complete transparency and reproducibility, through them users can execute their studies on a number of different platforms and systems (e.g. HPC system, cloud services, PC, etc.). Just like the QuNex source code, QuNex containers are also completely free and open to the research community.

### Containerization and Deployment

Through containerization, QuNex is fully platform-agnostic and comes in the form of both Docker and Singularity containers. This offer several advantages to end users. First, the QuNex container includes all of the required dependencies, packages and libraries which greatly reduces the time a user needs to setup everything and start processing. Second, the QuNex container is meticulously versioned and archived, which guarantees complete reproducibility of methods. Last but not least, containers can be run on practically every modern operating system (e.g. Windows, macOS, Linux) and can be deployed on any hardware configuration (e.g. desktop computer, laptop, cloud, high performance computing system). Users can easily execute the QuNex container via the included `qunex_container` script, which removes common technical barriers to connecting a container with the operating system. Furthermore, when running studies on an HPC system users need to manually configure the parameters of the underlying scheduling system, which can be again a tedious task for those that are not familiar with scheduling system. To alleviate this issue, the `qunex_container` script offers native support for several popular job schedulers (SLURM, PBS, LSF).

### QuNex Commands

A detailed list and a short description of all commands, along with a visualization of how commands can be chained together, can be found in the **Supplementary Information**. Here, we specify a short description for each of the functional groups of QuNex commands.

#### Study creation, data onboarding and mapping

This group of commands serves for setting up a QuNex study and its folder structure, importing your data into the study and preparing all the support files required for processing.

#### HCP Pipelines

These commands incorporate everything required for executing the whole HCP MPP along with some additional HCP Pipelines commands. Commands support the whole HCP MPP along with some additional processing and denoising commands. Below is a very brief overview of each pipeline, for details please consult the manuscript prepared by Glasser et al. (9) and the official HCP Pipelines repository (https://github.com/Washington-University/HCPpipelines). See **Figure S3** for a visualization of HCP Pipelines implementation in QuNex.

#### Quality control

QuNex contains commands through which users can execute visual QC for a number of commonly used MRI modalities – raw NIfTI, T1w, T2w, myelin, fMRI, dMRI, eddyQC, etc.

#### Diffusion analyses

QuNex also includes functionality for processing images acquired through dMRI. These commands prepare the data for a number of common dMRI analyses including diffusion tensor imaging (DTI) and probabilistic tractography.

#### BOLD analyses

Before running task-evoked and restingstate functional connectivity analyses, BOLD data needs to be additionally preprocessed. First, all the relevant data needs to be prepared – BOLD brain masks need to be created, BOLD image statistics need to be computed and processed and nuisance signals need to be extracted. These data are then used to process the images, which might include spatial smoothing, temporal high and/or low pass filtering, assumed HRF and unassumed HRF task modeling and regression of undesired nuisance and task signal.

#### Permutation Analysis of Linear Models (PALM)

The main purpose of this group of commands is to allow easier use of results and outputs generated by QuNex in various PALM (12) analyses (e.g. second-level statistical analysis and various types of statistical tests).

#### Mice pipelines

QuNex contains a set of commands for onboarding and preprocessing rodent MRI data (typically in the Bruker format). Results of the mice preprocessing pipelines can be then analysed using the same set of commands as with human data.

