## Supplementary Materials for "QuNex – An Integrative Platform for Reproducible Neuroimaging Analytics"

### Supplementary Information

**Validation Datasets.** The datasets used to test preprocessing via QuNex are described briefly below.

Public datasets:

1. ABIDE-I and II. Autism Brain Imaging Data Exchange I and II (65, 66). NDA Collection ID: 2039. International consortium focused on sharing data to inform neural mechanisms underlying autism spectrum disorder (ASD). ABIDE-I includes structural MRI, resting-state fMRI, and phenotypic data from 1112 subjects (539 from individuals with ASD and 573 from typical controls, ages 7-64 years, median 14.7 years across groups) across 17 sites. ABIDE-II includes data from 19 sites totalling 1114 subjects (521 individuals with ASD and 593 controls, age range: 5-64 years).
2. B-SNIP I. Bipolar-Schizophrenia Network on Intermediate Phenotypes (67). NDA Collection ID: 2274. Consortium with 6 sites across North America focused on trans-diagnostic psychosis spectrum disorders (schizophrenia, bipolar disorder with psychotic features, schizoaffective disorder with psychotic features). Data from patient first-degree relatives and healthy controls were also collected.
3. EMBARC. Establishing Moderators/Biosignatures of Antidepressant Response - Clinical Care study (68). NDA Collection ID: 2199. EMBARC is a clinical trial designed to compare three mechanistically distinct treatments for Major Depressive Disorder (MDD) (citalopram, bupropion, and cognitive behavioral therapy).
4. HCP. Human Connectome Project Young Adult cohort (21). NDA Collection ID: 2825. Data collection of high-resolution imaging and phenotype data totalling 1,200 normal young adults, aged 22-35.
5. PNC. Philadelphia Neurodevelopmental Cohort (69). dbGaP Study Accession: phs000607.v3.p2. Large study with diverse cohort of >9500 youths ages 8–21 years old from the greater Philadelphia area who received medical care at the Children's Hospital of Philadelphia. A subset of these individuals (n=1445) underwent neuroimaging.
6. NAPLS3. North American Prodrome Longitudinal Study (70). NDA Collection ID: 2275. Longitudinal study of 378 individuals at clinical high risk (CHR) of psychosis and 162 matched controls across 9 North American sites.
7. CamCAN. Cambridge Centre for Ageing and Neuroscience (Cam-CAN). `www.cam-can.org`. Collaborative study investigating the effects of ageing and cognition across the lifespan. Includes data from individuals ages 18 and up.

Internal datasets:

1. Lj-Clin. Combined data from two ongoing clinical studies collected at the University of Ljubljana, including: Subjective cognitive complaint (n=26); Transcranial magnetic stimulation in MDD (n=54).
2. Lj-Long. Composite data from 4 longitudinal datasets collected at the University of Ljubljana, including: Neurovacation (n=36 first visits, n=30 follow-up sessions); Neurovacation-2 (n=21 first sessions with n=19 follow-up sessions); BW-L, (n=39 first sessions with n=31 follow-up sessions); MMI (n=45 with 1-3 sessions).
3. Lj-Pharma. Pharmacological dataset of hyperglycaemia collected at the University of Ljubljana. N=41 participants (kids mean age 14y) with two sessions each participant.
4. Lj-Basic. Composite data from 14 smaller studies of cognitive functions in healthy young adults collected at the University of Ljubljana. These studies incldued: "attentional blink" (n=31); Oddball (n=34); WM-1 (n=32); WM-2 (n=32); WM-3 (n=35); sWM (n=31); sWM-D (n=28); sWM-UI (n=33); sWM-UIR (n=30); NEco (n=39); BW (n=47); C3T-1 (n=51); C3T-2 (n=49); LOC (n=24).
5. Yale-Clin. Clinical data collected on a Siemens 3T Prisma scanner at Yale University, including 30 patients with schizophrenia, 39 patients with obsessive compulsive disorder, and 36 matched controls.
6. Yale-Ket. Pharmacological challenge study collected at Yale University. Included scans from 40 healthy adults who underwent acute ketamine infusion during the scan, and 40 patients with schizophrenia.
7. Zurich-Psi. Pharmacological challenge study with psilocybin collected at University of Zurich (71). Includes 23 healthy adults who each underwent 2 sessions.
8. Zurich-LSD. Pharmacological challenge study with LSD collected at University of Zurich (72). Includes 24 healthy adults who each underwent 3 scans.

DRAFT

**Supported External Tools and Pipelines.** QuNex is designed to maximize interoperability between community-developed neuroimaging tools. Below is a list of tools, software, and pipelines which are currently integrated into the QuNex platform, as well as those which are planned for future integration (indicated by *).

- dicomifier* (73)
- dcm2niix (74)
- dicm2nii (75)
- ANTs* (76)
- AFNI* (5)
- SPM* (8)
- FSL (7), including FreeSurfer (6); PALM (12)
- HCP MPP (9)
- HCP Workbench (21)
- HCP ASL (arterial spin labelling) (77)
- fMRIPrep* (10)
- Mouse resting-state fMRI pipelines (19)

**Descriptions of Individual QuNex Commands.** In this section, we provide an overview and description of the QuNex commands currently supported. For a visual representation of commands supported in QuNex see **Figure S1**.

##### *Study Creation, Data Onboarding, and Mapping.*

**create_study** creates the base study folder structure (**Figure S2**). Inside the folder structure QuNex will store everything that is relevant for your work. The intention of the `info` subfolder is to store basic study information. In the `sessions` subfolder, QuNex will store all session data. The `processing` subfolder will contain processing logs, lists and scripts, while the `analysis` subfolder will store analysis scripts and group level results.

**import_dicom** is used to simplify and streamline importing of DICOM files into your QuNex study. The command automatically processes packets with individual MRI sessions's DICOM or PAR/REC files and generates NIfTI files that will be later used in processing.

**import_bids** command serves a similar purpose as `import_dicom`, it processes a BIDS formated dataset and imports the data into a QuNex study folder.

**import_hcp** is used to import dataset strucured as Human Connectome Project data (HCP lifespan and HCP young adults) into a QuNex study folder.

**bruker_to_dicom** this command can be used to convert data in the Bruker file format (a format commonly used when acquiring rodent MRI data) to DICOM. Once converted, the dataset can be onboarded into QuNex using the `import_dicom` command.

**create_session_info** creates session files that hold the information about MRI recordings of each session. These files are necessary for correct mapping to a folder structure supporting specific pipeline processing.

**create_batch** creates a joint batch file from all the individual session files. The batch file thus contains concatenated information about all the sessions of a study. Besides session information batch file also stores values for processing parameters that are valid for all the sessions. You can use the created batch file to manually tune these parameters.

**setup_hcp** maps images to a folder structure that conforms to the naming conventions used by the HCP MPP workflow to enable HCP MPP processing.

##### *HCP Pipelines.*

**hcp_pre_freesurfer** runs the PreFreeSurfer pipeline of the HCP MPP. The command averages any image repeats (i.e. multiple T1w or T2w images available), creates a native, undistorted structural volume space for the subject, provides an initial robust brain extraction, aligns the T1w and T2w structural images (register them to the native space), performs bias field correction and registers the subject's native space to the MNI space.

**hcp_freesurfer** runs the FreeSurfer pipeline of the HCP MPP. The HCP FreeSurfer pipeline performs spline interpolated downsampling, recon-all (a robust pipeline optimized for 1 mm isotropic data (78–80), in this and HCP Pipeline's case optimized for the higher resolution HCP Phase II data), high-resolution white matter placement, fine tuning of the T2w to T1w registration and high-resolution pial tuning.

**hcp_post_freesurfer** runs the PostFreeSurfer pipeline of the HCP MPP. This pipeline performs the conversion of FreeSurfer volumes and surfaces to CIFTI format, creates the FreeSurfer ribbon file at full resolution and performs the myelin mapping.

**hcp_fmri_volume** runs the fMRIVolume pipeline of the HCP MPP. This pipeline is used to perform the following operations on BOLD image files: slice timing correction, gradient distortion correction, motion correction, echo-planar imaging distortion correction, registration to the T1w image, one step resampling to target space, intensity normalization and bias removal.

**hcp_fmri_surface** runs the fMRISurface pipeline of the HCP MPP, which maps the BOLD signal from cortical gray matter to surface representation and creates the fMRI ribbon, smooths the surface, executes subcortical processing and generates dense timeseries in CIFTI format.

**hcp_diffusion** runs the Diffusion pipeline of the HCP MPP. Note that for this pipeline the dMRI data should be acquired in pairs of phase encoding reversed images. This pipeline starts by intensity normalizing the mean $b_0$ image across the six diffusion series. After the $b_0$ intensity normalization, the $b_0$ images of both phase encoding directions are used to calculate the susceptibility-induced $B_0$ field deviations. Next, the full timeseries from both phase encoding directions is used in the FSL's "eddy" tool (81, 82) for modeling of eddy current distortions and subject motion. Gradient distortion is corrected and the $b_0$ image is registered to the T1w image using FLIRT's boundary-based registration (BBR) cost function (83). The diffusion data output from eddy are then resampled into 1.25 mm native structural space and masked. Diffusion directions and the gradient deviation estimates are also appropriately rotated and registered into structural space.

**hcp_asl** runs the HCP ASL (arterial spin labeling) pipeline. Note here that this pipeline is not part of the official HCP Pipelines repository and is developed and maintained by a research group at the University of Nottingham (`https://github.com/physimals/hcp-asl`).

##### *Additional HCP Pipelines Commands.*

**hcp_icafix** runs the ICAFix pipeline of the HCP Pipelines. HCP ICAFix first performs the spatial ICA decomposition of the data and then attempts to auto-classify ICA components into signal and noise. Noise components are then removed from the 4D fMRI data. If `hcp_icafix`

DRAFT

command finishes successfully the hcp_post_fix command is ran automatically by default.

**hcp_post_fix** runs the PostFix pipeline of the HCP Pipeline which creates Connectome Workbench scene files that can be used to visually review the signal versus noise classification generated by HCP ICAFix.

**hcp_reapply_fix** command combines two HCP Pipelines that are almost always run in sequence. First, it runs the ReApplyFix HCP Pipeline which applies the hand reclassifications of noise and signal components from ICAFix using the ReclassifyAsNoise.txt and ReclassifyAsSignal.txt input files. Next, it executes the HCP Pipeline's ReApplyFix which reapplies Fix cleanup to the volume and the default CIFTI.

**hcp_msmall** runs the MSMAll HCP Pipeline which takes care of surface-based functional alignment. It computes the MSMAll registration based on resting-state connectivity, resting-state topography and myelin-map architecture. HCP MSMAll should only be used on ICAFix cleaned data. After a successful hcp_msmall run hcp_dedrift_and_resample will be run automatically by default.

**hcp_dedrift_and_resample** runs the DeDriftAndResample HCP Pipeline which is used to resample the data according to the MSMAll registration.

**hcp_make_average_dataset** runs the make average dataset command from the HCP Pipelines. The command makes an average dataset across multiple sessions or subjects that is required to run the hcp_temporal_ica command.

**hcp_temporal_ica** runs the temporal ICA pipeline of the HCP Pipelines. This commands performs temporal ICA decomposition of the data and then attempts to auto-classify the independent components into signal and noise. The hcp_make_average_dataset commands needs to be ran before hcp_temporal_ica when using multiple sessions or subjects.

**map_hcp_data** . Finally, once images are preprocessed using the HCP preprocessing pipelines and verified using the QC tools, the results need to be mapped back to the QuNex folder structure. This can be accomplished using the map_hcp_data command.

***Quality Control.***

**run_qc** runs the quality control (QC) for a specified modality or a processing command. Through this function QuNex is capable of performing raw NIfTI, T1w, T2w, myelin, fMRI, dMRI, eddyQC and general QC. The command is compliant with both legacy data (without T2w scans) and HCP-compliant data (with T2w and dMRI scans).

**dwi_eddy_qc** command is based on FSL's eddy to perform quality control on diffusion MRI (dMRI) data. It explicitly assumes that eddy was ran (eddy is ran in the hcp_freesurfer command).

***Diffusion Analyses.***

**dwi_dtifit** carries out diffusion tensor fitting at each voxel using FSL's DTIFIT. Its outputs include mean diffusivity and fractional anisotropy. Note that dwi_dtifit is not required for probabilistic tractography. HCP's diffusion results must be present before running this command.

**dwi_bedpostx_gpu** models crossing fibers within each voxel using FSL's BEDPOSTX. The probability of diffusion from each voxel in every direction versus all other directions is estimated, building the distributions that are necessary for running probabilistic tractography. HCP MPP diffusion results must be present before running the command. This command requires a GPU device for successful execution.

**dwi_xtract** executes the FSL's XTRACT (cross-species tractography) command (14). It can be used to automatically extract a set of carefully dissected tracts in humans and macaques. It can also be used to define one's own tractography protocols where all the user needs to do is to define a set of masks in standard space (e.g. MNI152).

**dwi_pre_tractography** runs whole-brain (dense) pre-tractography trajectory space generation and is a prerequisite for dwi_probtracx_dense_gpu.

**dwi_probtracx_dense_gpu** samples the BEDPOSTX distribution results using FSL's PROBTRACKX. A whole-brain dense connectome is generated showing the probability of streamline connections from every voxel to every other voxel. The command can produce two versions of the dense connectivity matrix based on different seeding strategies, Matrix 1 computes probabilistic tractography between each grey matter point and every other grey matter point, while Matrix 3 starts with a white matter voxel and computes the tractography in both directions (along a given orientation) to the two grey matter points at either end. Whereas Matrix 1 is unidirectional surface-to-surface, Matrix 3 is bidirectional voxel-to-surface which better reflects long-range projections. This command requires a GPU device for successful execution.

**dwi_seed_tractography_dense** reduces the dense connectome using a given "seed" structure (e.g. thalamus). The produced outputs will contain streamline information for only those connections originating from the specific anatomical seed.

***BOLD Analyses.***

**create_bold_brain_masks** command creates, for each BOLD separately, a mask of brain voxels for which signal is present. This ensures that in the following commands neither the signal outside of the brain, nor the areas within the brain from which signal was not captured due to small field of view or subject movement between the BOLD image acquisitions, are used. For instance, due to small field of view, inferior parts of cerebellum might not have been imaged and the signal is missing.

DRAFT

**compute_bold_stats** computes a number of frame-by-frame statistics for BOLD timeseries including signal change between successive frames. These statistics allow assessment of the BOLD timeseries quality and identification of frames with artifact in the signal.

**create_stats_report** commands creates and evaluates reports based on statistics computed in the `compute_bold_stats` command. Reports are prepared across BOLD images for each individual and compiled into a joint group report. The command also evaluates movement and intensity changes statistics according to provided criteria and identifies frames that might have movement artifacts (i.e., "bad frames") to allow their exclusion or interpolation in commands that follow.

**extract_nuisance_signal** command is used for estimating nuisance (and potentially other) signals from the brain. These signals are usually comprised of mean signals from brain ventricles, white matter and gray matter. This command allows extraction of these signals – as well as signals from any additional specified region or regions – from the brain to be used in the analyses that follow.

**parcellate_bold** command is used for parcellation of the BOLD dense files using whole-brain parcellations (e.g. the Glasser parcellation with subcortical labels).

**create_conc** command can be used to generate concatenation (conc) files. Conc files are text files that offer a simple way of specifying a list of imaging files that should be concatenated during various processing and analysis commands.

**join_fidl** command creates a joint fidl file – a text file that provides the information on events during the task BOLD runs – based on separate fidl files for each functional image and a related conc file. For example, when fidl files were generated for each BOLD image separately, this command enables combining of fidl files for all the relevant BOLD images into a single fidl file. When combining the fidl files the times are recomputed to correctly match the concatenated BOLD signal timeseries.

**check_fidl** command creates a visual overview of the events and their durations based on the provided fidl file. Specifically, it creates a plot of events and their durations and saves them in PDF files for quick visual review.

**preprocess_bold** command performs the actual processing of the data. This can include spatial smoothing, temporal high and/or low pass filtering, task GLM modeling and nuisance signal regression in any desired sequence and combination. The results are GLM beta estimates, which can be used in task activation analyses or other purposes and preprocessed residuals after removal of undesired signal, which can be used as the input for functional connectivity analyses. The `preprocess_bold` command is used when each BOLD image is processed independently, which is most relevant for resting state data or tasks with a single BOLD run acquisition.

**preprocess_conc** this command has the same functionality as `preprocess_bold` and is used in the case when multiple BOLD images are to be concatenated and processed jointly (as is the case in task designs that span multiple BOLD images).

**create_list** command can be used to generate list ("list") files. List files are used to specify related files of interest of different types across a group of sessions, e.g. roi, fidl, conc, bold files for each session. List files allow enumerating all the relevant files needed to perform specific analyses, e.g. computation of seed-based functional connectivity, on a group of sessions (or subjects).

**general_extract_roi_glm_values** command is used to extract mean activation (beta) estimates of specified effects for the specified ROIs from volume or CIFTI GLM files. The resulting beta estimates or percent signal change values can be saved in simple text files in either long or wide format so that they can be easily imported in other tools for further processing, analysis and visualization.

**general_extract_glm_volumes** command extracts GLM estimates of the effects of interests, next it organizes and saves these estimates in the desired manner (specific order within a single file or into individual files). This step enables desired organization of imaging data for running second-level analysis in PALM.

**fc_compute_roifc** computes ROI-based functional connectivity matrices for an individual subject/session or a group of sessions.

**fc_compute_seedmaps** computes seed based functional connectivity maps for individual subject/session or a group of sessions.

**fc_compute_gbc** commands are used for computing global brain connectivity (GBC). There are two command for this purpose. The `fc_compute_gbc3` can be used for computing GBC maps for individuals as well as group. `fc_compute_gbcd` computes GBC averages for each specified ROI for $n$ bands defined as distance from ROI.

***Permutation Analysis of Linear Models (PALM).***

**create_ws_palm_design** command can be used to prepare PALM design specification files. The files are used later in the PALM analysis and include information about the design matrix, t-contrasts, f-contrasts and exchangeability blocks, respectively.

**run_palm** command runs second level analysis using PALM permutation resampling. It can be run on grayordinate, CIFTI dense or parcellated images or volume, NIfTI images. The output of this command is a number of different image files that contain results for all t- or f-contrasts provided by the "design" parameter and different methods for multiple testing correction.

DRAFT

**mask_map** command enables easy masking of CIFTI images (e.g. ztstat image from PALM), using the provided list of mask files (e.g. p-values images from PALM) and thresholds. More than one mask can be used in which case they can be combined using a logical OR or AND operator.

**join_maps** command concatenates the listed CIFTI maps and names the individual volumes, if names are provided.

**general_find_peaks** performs image smoothing and identifies peak ROI using a watershed algorithm to grow regions from identified peaks.

***Non-human Primates Processing.*** Current stable and publicly released version of QuNex incorporates processing commands required to perform post-mortem macaque tractography. Some steps used in this pipeline (`create_study`, `dwi_dtifit`, `dwi_bedpostx_gpu`, `fsl_xtract`) are already described in previous sections.

**import_nhp** similar to other import commands in this case enabling onboarding of post-mortem macaque data into a QuNex study.

**dwi_f99** executes the FSL's F99 script for registering own diffusion or structural data to the F99 atlas. This atlas is used when processing macaque data.

***Mice Pipelines.***

**setup_mice** runs the command to prepare a QuNex study for mice preprocessing. The command can perform the following steps if they are necessary: voxel size increase, orientation correction, TR unit correction and despiking.

**preprocess_mice** tuns the QuNex mice preprocessing command. The command first performs MELODIC (Multivariate Exploratory Linear Optimized Decomposition into Independent Components), which is followed by FSL's FIX step. After these two steps the command generates a QC image. Finally, bandpassed data is mapped to both EPI and ABI (Allen) templates.

**map_mice_data** . Once images are preprocessed using the mice preprocessing pipelines, the results need to be mapped back to the QuNex folder structure in order to use them in the following analyses. This can be accomplished using the `map_mice_data` command.

**Visual Quality Control (QC).** Visual inspection of input data and QuNex outputs is a critical step in ensuring that data are accurately preprocessed and of sufficient quality for further analytics. QuNex includes native functionality via `run_qc` for generating scenes and PNG images for rapid inspection of data and preprocessing results. These images are generated at the level of individual scans and are supported for multiple modalities, including T1w, T2w, myelin, fMRI, and dMRI (**Figure S4A**). This allows users to visually check both the quality of their input scans and the outputs of intermediate steps, including FreeSurfer segmentation and surface tracing for structural data, motion and signal-to-noise (SNR) of functional scans, and calculation of diffusion metrics for dMRI data. Additionally, QuNex also generates motion plots for individual fMRI scans (**Figure S4B**). This includes the estimated motion parameters (dx, dy, dz and x, y, z) for every frame in each functional scan (left), as well as timeseries "greyplots" and metrics for motion scrubbing (frame displacement/FD, image intensity normalized root mean squared error (RMSE)/dvarsm (84)). This allows users to easily identify sessions/scans with low versus high motion. The generation of lightweight files (PNG, PDF) for visual QC is designed to allow users to quickly survey data quality for large numbers of scans.

**QuNex Parallelism and Scheduler Support.** Neuroimaging studies contain large amounts of acquired data that needs to be processed and analysed. Doing so requires a lot of processing power and is traditionally executed on HPC systems in a parallel fashion. To enable parallel processing QuNex commands accept three different parameters that determine the parallel processing regime. The `parjobs` parameter determines the maximum number of independent jobs that will be scheduled for execution on the HPC system. The `parsessions` parameter determines how many sessions at most will be ran in parallel inside each individual job. Finally, the `parelements` parameter determines the number of elements that will be processed in paralell inside each session. For example in `hcp_fmri_volume` and `hcp_fmri_surface` the `parelements` parameter determines how many BOLDs will be processed in parallel per each session. See **Figure S7** for a visual representation of the logic behind the QuNex's parallelism parameters.

On HPC systems users do not execute commands in the same fashion as on their own PCs, instead additional work is needed. In order to execute commands on HPC system, commands need to be scheduled via specialised scheduling systems, which is usually done via scheduling scripts. Furthermore, format and scheduling script parameters are scheduler dependant, for example a script for the SLURM scheduling system would be different than the one for PBS or LSF. **Figure S5** visualizes an example of such a script for the SLURM scheduling system.

To make QuNex as user friendly as possible we implemented native support for three popular scheduling systems: SLURM, PBS and LSF. The only change needed to schedule a QuNex command as a job on an HPC system is to add a `scheduler` parameter when executing commands. QuNex will automatically prepare the scheduling script and send it for execution to the scheduling system. This saves users work, particularly when users want to split their execution into several jobs (as often done to optimize performance). Without QuNex's native scheduling support, users would have to write up a scheduling script for each individual job. The figure below (**Figure S6**) shows an example QuNex call that facilitates the `scheduler` parameter. The example on the figure will schedule a job for each of the sessions in the study. The number of jobs that will run in parallel depends on the study size and the capabilities of your computation

DRAFT

system. Within each session, 2 BOLDs will be processed simultaneously. Without the `scheduler` this would require multiple scheduling scripts, one for each job. Furthermore, users can use the `parjobs`, `parsessions` and `parelements` parameters to fine tune processing to their needs. QuNex will automatically reserve processing resources via the scheduling system's parameters in an efficient way. For example, when using SLURM, QuNex will automatically set the `cpus-per-task` parameters. This automatic setup of scheduling parameters should ensure a decent processing performance in the vast majority of use cases. If users are not satisfied with the automatic configuration they can easily override by manually setting relevant scheduling system's parameters through the `scheduler` parameter.

**XNAT Deployment.** XNAT, an open source informatics platform, is widely used as a neuroimaging project management and data storage system (33). QuNex is fully interoperable with XNAT and can deploy preprocessing and analytic workflows natively on XNAT. QuNex is containerized and deployed in XNAT via a container plugin. XNAT provides a frontend UI for easy access to data, as well as a raw DICOM web viewer (**Figure S8**). The QuNex container is pulled into XNAT from the container repository, at which point the container specification can be created. The container specification lays out what processing steps are to be used. This is the equivalent of a command line argument. Several commands / specifications can be created to granularize what QuNex commands to run. Sessions can be launched individually, or multiple sessions can be run in parallel via XNAT's processing dashboard. After the container is launched, the user can view the StdOut in realtime to track job status. Depending on what commands were run, once the container finishes, QC images are available to view from the web browser. Such interoperability greatly streamlines neuroimaging workflows and faciliates data access and sharing.

**Interoperability with Analytical Tools and Independent Features.** The outputs of QuNex are desigend to be interoperable with further analytics and other analytical tools. Neuroimaging files can be manipulated via NIFTI or CIFTI-compatible tools, such as FSL, AFNI, HCP Workbench, nipy etc; users can also specify, for many QuNex feature-generating commands such as `fc_compute_seedmaps`, that results are additionally output as plain numerical tables that can be imported into any data manipulation software. The surface-based cortical representation allows comparison with independent molecularly-informed neural features, such as gene expression maps (85, 86). Here we highlight previously published studies which have leveraged QuNex workflows.

QuNex is capable of processing and analysis of pharmacological neuroimaging datasets designed to investigate the effect of pharmacological manipulation on neuroimaging features. In one exemplar study (72), subjects each underwent fMRI scanning under 3 counterbalanced and blinded conditions: 1) two doses of placebo pills (Placebo condition); 2) placebo then LSD (lysergic diethylamide acid; LSD condition); 3) pretreatment by ketanserin followed by LSD (Ket + LSD condition). Ketanserin is a select $5-HT_{2A}$ receptor antagonist whereas LSD has a diffuse profile of receptor targets, including $5-HT_{2A}$, $_{2C}$, $_{1A/B}$, $_{6}$, and $_{7}$ as well as dopamine D1 and D2 receptors. The study was designed to test the hypothesis that the neural effects of LSD are largely modulated by the 5-HT2A receptor; thus, pretreatment by ketanserin should block the effect of LSD, resulting in a neural map that was highly similar to that of the Placebo condition (**Figure S9A**). The differential global brain connectivity (ΔGBC) results between LSD and (Ket + LSD) conditions indeed showed that the within-subject contrast of LSD versus (Ket + LSD) conditions was highly spatially correlated (r=0.88) with the LSD versus Placebo contrast. This strongly implicates the role of $5-HT_{2A}$ receptors in governing the neural effect of LSD. Such neural features and statistical maps are easily generated by the preprocessing and analytic mechanisms enabled by QuNex.

QuNex outputs can also be integrated with behavioral phentoype data. In a recent study (50), dimensionality reduced axes of symptom variation were mapped to GBC across 436 patients with a psychosis spectrum disorder (PSD, **Figure S9B**). The principal components (PC) of symptom variation (here PC3 is shown as an exemplar) were oblique to traditional symptom factors [Positive, Negative and General factors from the Positive and Negative Syndrome Scale (PANSS) and Cognitive factor from the Brief Assessment of Cognition in Schizophrenia (BACS)] and comprised a linear combination of the individual items on these assessments. Importantly, PC scores did not differ at the mean level between diagnostic groups (CON, control; PSD; all PSD patients; BPP, bipolar disorder with psychotic features; SADP, schizoaffective disorder with psychotic features; SZP, schizophrenia) but the coefficient map of PC3 scores regressed on to GBC was statistically stronger than both the coefficient maps of Positive and Negative symptom scores.

Data processed via QuNex can be seamlessly integrated with other surface-based neural features, such as cortical gene expression from the Allen Human Brain Atlas (AHBA) (85), as shwon in **Figure S9C**. The process of mapping AHBA gene expression patterns to derive group-average parcellated cortical surface expression maps, per (86), is briefly outlined here. Genes expression levels measured with DNA microarray probes were sampled from hundreds of neuroanatomical structures in the left hemisphere of six postmortem human brains. At the single-subject level, samples for each subject were first mapped from the volumetric space onto that subject's native reconstructed two-dimensional cortical surface. Next, parcellated gene expression maps were constructed for each subject, by interpolating the expression of each gene to fill parcels from the HCP multimodal parcellation (MMP1.0) (23). Finally, a group-level parcellated expression map for each unique gene was computed by averaging parcellated expression levels across subjects' selected gene probes. For details refer to (86). Gene expression maps can be used as an independent source of molecular information for anchoring fMRI-derived neuroimaging maps, such

DRAFT

as the LSD data shown above (72). The Z-scored contrast of LSD > placebo in N=24 subjects is shown, highlighting elevated GBC (GBC) (87) under LSD in sensory areas such as primary visual and somatosensory cortices and reduced GBC in areas such as retrospenial cortex. The receptors implicated in LSD pharmacology are differentially expressed across the cortex. To investigate LSD's receptor pharmacology, we correlated the LSD > Placebo map with the gene expression map of HTR2A, resulting in a correlation of $r_p = 0.50$, $p < 0.001$. The correlation between the LSD > Placebo map and HTR2A expression was higher than with any other receptor gene of interest (DRD1, DRD2, HTR1A, HTR2C, and HTR7), and was in fact higher than 95.9% of all possible correlations across all 20,737 available genes. These results show that LSD-induced changes in functional connectivity quantitatively match the spatial expression profile of genes coding for the 5-HT2A receptor, further supporting the central role of this receptor system in LSD's effects.

**A Visualization of the QuNex Extensions Framework.** QuNex is completely open source and also designed with community contributions in mind. At the core of this is the QuNex extensions framework, which allows researchers to develop their own modules and seamlessly attach them to the QuNex platform. By doing so these extensions become interoperable and can facilitate several functionalities already implemented into QuNex (e.g., scheduling, parallelism, logging, etc.). A visualization of the QuNex extensions framework is shown in **Figure S10**.

DRAFT

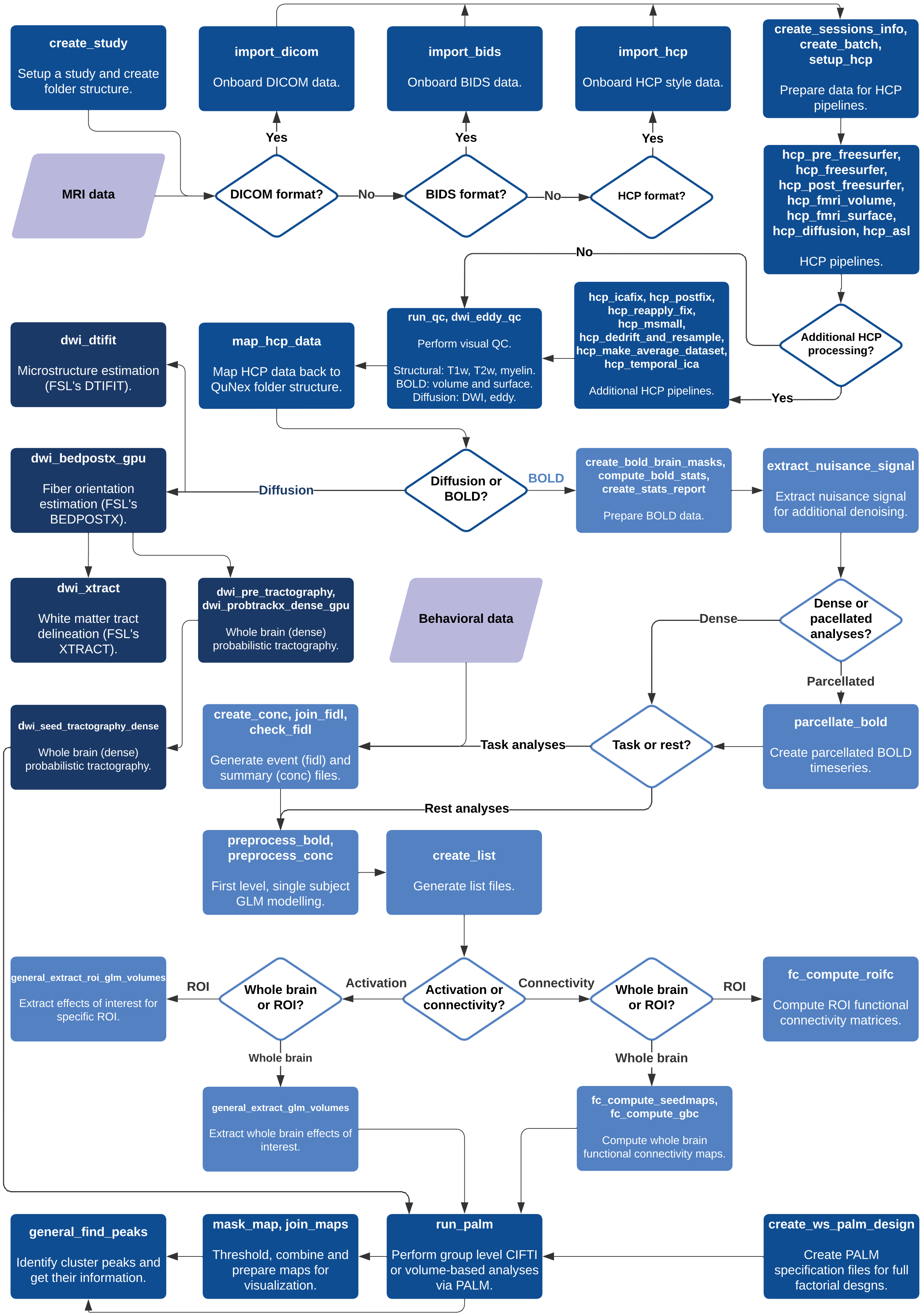


**Fig. S1. A Visual Representation of QuNex Commands and Their Dependencies.** QuNex Commands are visualized with colored rectangles, color coding denotes groups of commands (e.g., onboarding, preprocessing, dMRI, fMRI, analyses, etc.). Data inputs are visualized as parallelograms, while choices in processing and analyses (either researcher, data or study driven) are marked as rhombi. Arrows denote command dependencies. Note that only the commands that are primarily meant for processing and analysis of human data are visualized on this figure.

Example Folder Hierarchy Specification

```
Study                                 # -- Root folder of the whole study
│
├── analysis                          # -- Group analyses and results
│   └── scripts                       # -- Analyses scripts
│
├── info                              # -- Various information and materials
│   ├── bids                          # -- Study-level information in BIDS format
│   ├── demographics                  # -- Participant demographic information
│   ├── hcpls                         # -- Study-level information in HCPLS format
│   ├── stimuli                       # -- Task-related stimuli
│   └── tasks                         # -- Task-related information
│
├── processing                        # -- Processing information and materials
│   ├── lists                         # -- List files used in processing and analyses
│   ├── logs                          # -- Processing logs
│   │   ├── comlogs                   # -- Detailed output generated by executed commands
│   │   └── runlogs                   # -- General information for executed commands
│   ├── scenes                        # -- Misc scenes used in analyses and QC
│   │   └── QC                        # -- Additional custom QC scenes and associated files
│   │       ├── BOLD                  # -- BOLD QC scenes and associated files
│   │       ├── DWI                   # -- DWI QC scenes and associated files
│   │       ├── myelin                # -- Myelin QC scenes and associated files
│   │       ├── T1w                   # -- T1w QC scenes and associated files
│   │       └── T2w                   # -- T2w QC scenes and associated files
│   ├── scripts                       # -- Processing scripts
│   └── batch.txt                     # -- Batch files contain processing parameter, analysis parameters and sessions' basic information
│                                     # -- Batch files can be generated with the create_batch command
│
└── sessions                          # -- Data from study's sessions
    ├── archive                       # -- Backed up and archived raw data
    │   ├── behavior                  # -- Archived raw behavioral data
    │   ├── BIDS                      # -- Archived raw BIDS data
    │   ├── EEG                       # -- Archived raw EEG data
    │   ├── HCPLS                     # -- Archived raw HCPLS data
    │   └── MR                        # -- Archived raw MR data
    ├── inbox                         # -- Data used in processing and analyses
    │   ├── behavior                  # -- Behavioral data
    │   ├── BIDS                      # -- BIDS data
    │   ├── concs                     # -- Conc files
    │   ├── EEG                       # -- EEG data
    │   ├── events                    # -- Fidl files
    │   ├── HCPLS                     # -- HCPLS data
    │   └── MR                        # -- MR data
    ├── QC                            # -- Group-level quality control data for all sessions
    └── specs                         # -- Specification files to be used on the sessions
        └── batch_example.txt         # -- Example, template batch file used to specify processing and analysis parameters
```

**Fig. S2. An Example of the QuNex Study's Folder Structure Specification.** The logic behind the folder hierarchy is to provide a clear and predictable structure. That way, both users and tools can reference data in consistent locations within a unified naming grammar, which limits the need for referential databases and metadata management. The data hierarchy is defined primarily for individual sessions. Group-level data hierarchy specification (e.g. the analyses folder) is not fixed, thus enabling flexible internal organization according to specific needs and goals.

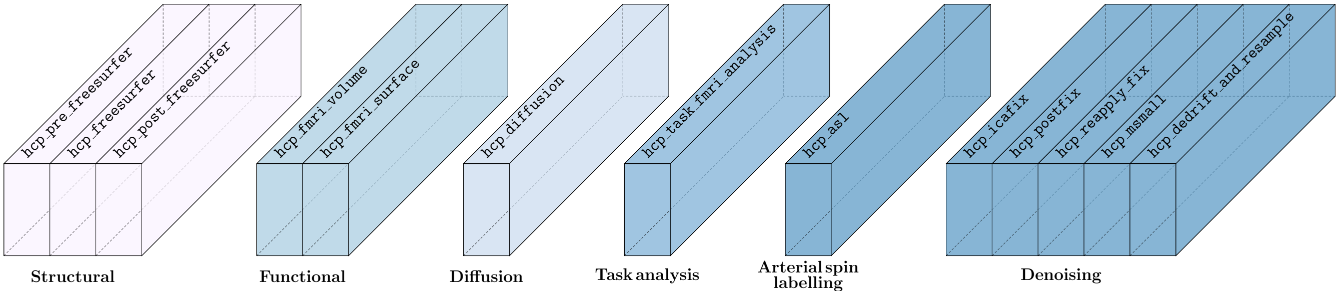


**Fig. S3. A Visual Representation of the HCP Pipelines implementation inside QuNex.** Labels on top of shapes denote QuNex command names. Labels below shape groups denote groups of commands. For example, to execute HCP ICAFix pipelines, one should run `qunex hcp_icafix`.

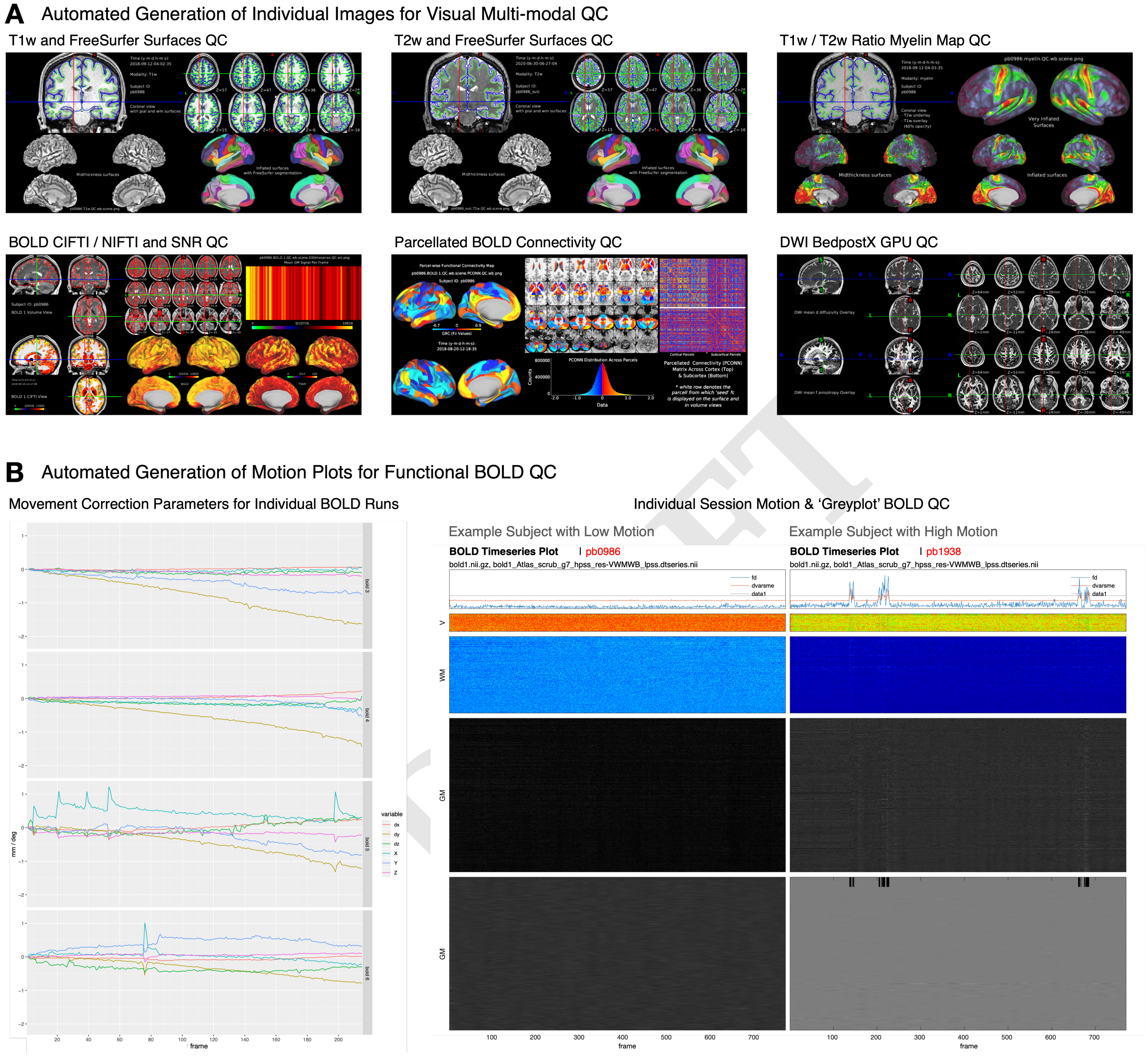


**Fig. S4. Automated visual QC engine for rapid inspection of data and outputs. A)** QuNex generates images for rapid and thorough inspection of data and processing outputs via the `run_qc` command. Images are generated for individual scans and allow the user to Supported modalities include NIFTI, T1w and T2w images with FreeSurfer segmentation results, T1w/T2w myelin maps, BOLD volume and surface mapping, parcellated BOLD data and functional connectivity results, and dMRI mean diffusivity and fractional anisotropy results. **B)** Motion parameters can be visualized for every frame in each functional BOLD scan. (Left) This includes the six motion paramters (dx, dy, dz and x, y, z) as well as metrics for motion scrubbing (frame displacement and image intensity normalized root mean squared error (RMSE)/dvarsm (84)). This allows users to quickly distinguish between subjects with low motion (middle panel) from those with high motion (right). Additionally, any specified nuisance regressors and the BOLD signal itself can also be visualized as greyplots to quickly evaluate the quality of each functional scan.

Example SLURM scheduling script

```
#!/bin/bash
#SBATCH --time=24:00:00
#SBATCH --ntasks=4
#SBATCH --cpus-per-task=5
#SBATCH --mem-per-cpu=4000
#SBATCH --partition=day
#SBATCH --job-name=hcp_volume

qunex hcp_fmri_volume \
    --sessions="/study/processing/batch.txt" \
    --sessionsfolder="/study/sessions" \
    --parsessions="4" \
    --parelements="5" \
    --overwrite="no"
```

**Fig. S5. An example of a SLURM scheduling script.** The above script would have to be prepared by users in order to schedule a run of QuNex command `hcp_fmri_volume`. The command would schedule a job where 4 sessions in parallel, within each session 5 BOLDs would be processed in parallel. Once such a script is prepared user can schedule it for execution via the SLURM's `sbatch` command.

DRAFT

Example QuNex call with scheduler

```
qunex hcp_fmri_volume \
    --sessions="/study/processing/batch.txt" \
    --sessionsfolder="/study/sessions" \
    --parelements="2" \
    --overwrite="no" \
    --scheduler="SLURM,time=24:00:00,mem-per-cpu=4000,partition
    =day"
```

**Fig. S6. Scheduling jobs directly via the QuNex command call.** By using the `scheduler` parameter one can prepare jobs for execution via the HPC's scheduling system in an easier and more compact fashion. Furthermore, it is possible to schedule multiple jobs at once where default values for reserving computational resources (number of CPUs per task) will be automatically set by QuNex depending on the amount of sessions and elements you want to run in parallel.

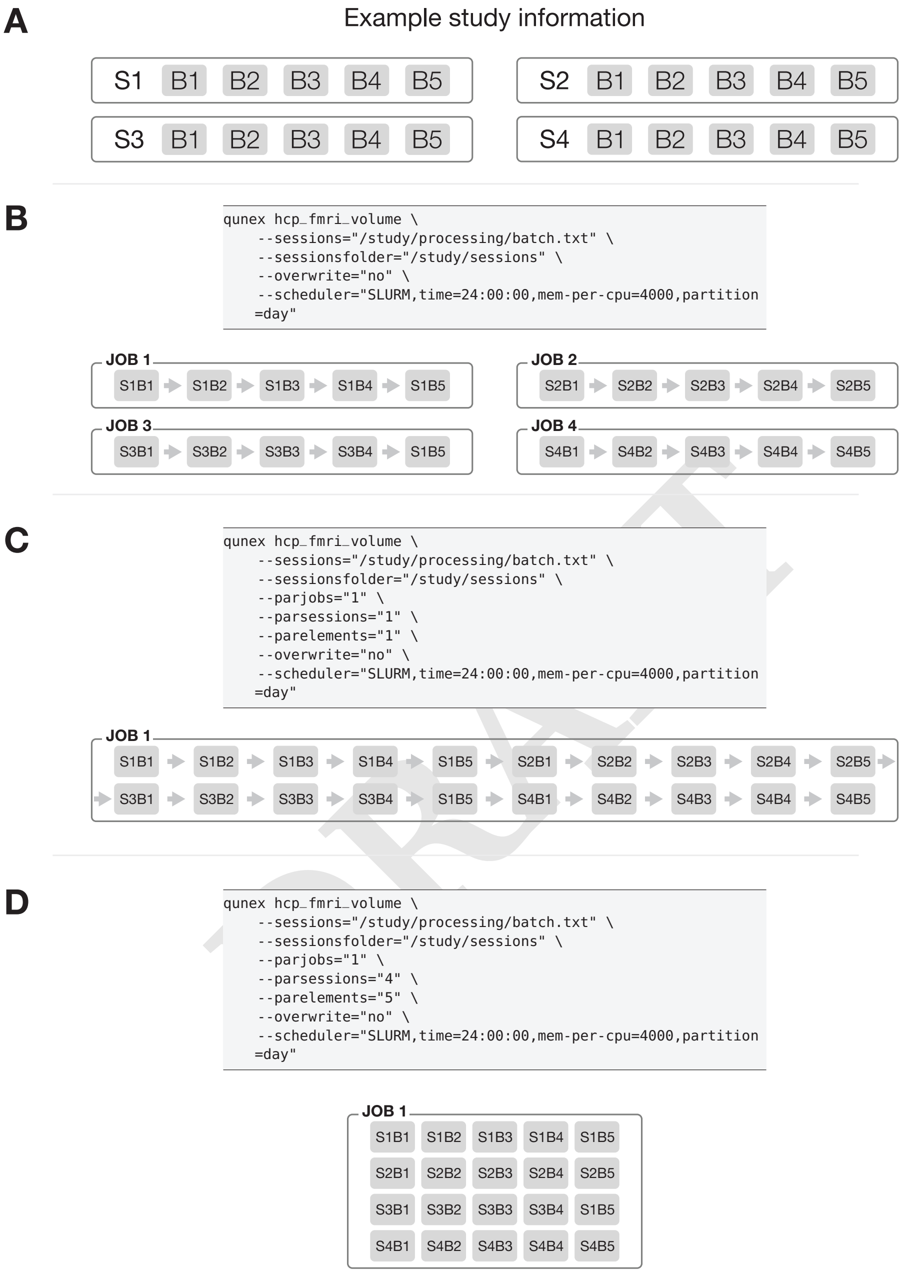


**Fig. S7. QuNex scheduling parameters and their effects. A)** Study information, our sample study has 4 sessions and each session has 5 BOLDs. Sessions are abbreviated with the letter S, while bolds are abbreviated with B. For example, B1 denotes BOLD 1 while S2B3 denotes session 2, BOLD 3. **B)** The default execution, QuNex will schedule an individual job for each session, within each sessions elements/BOLDs will run in a serial fashion unless the `parelements` parameter is also set. **C)** A completely serial execution – all elements/BOLDs will be executed in a serial fashion within a single job. **D)** Complete parallelism within a single job, where all elements/BOLDs will be executed in parallel.

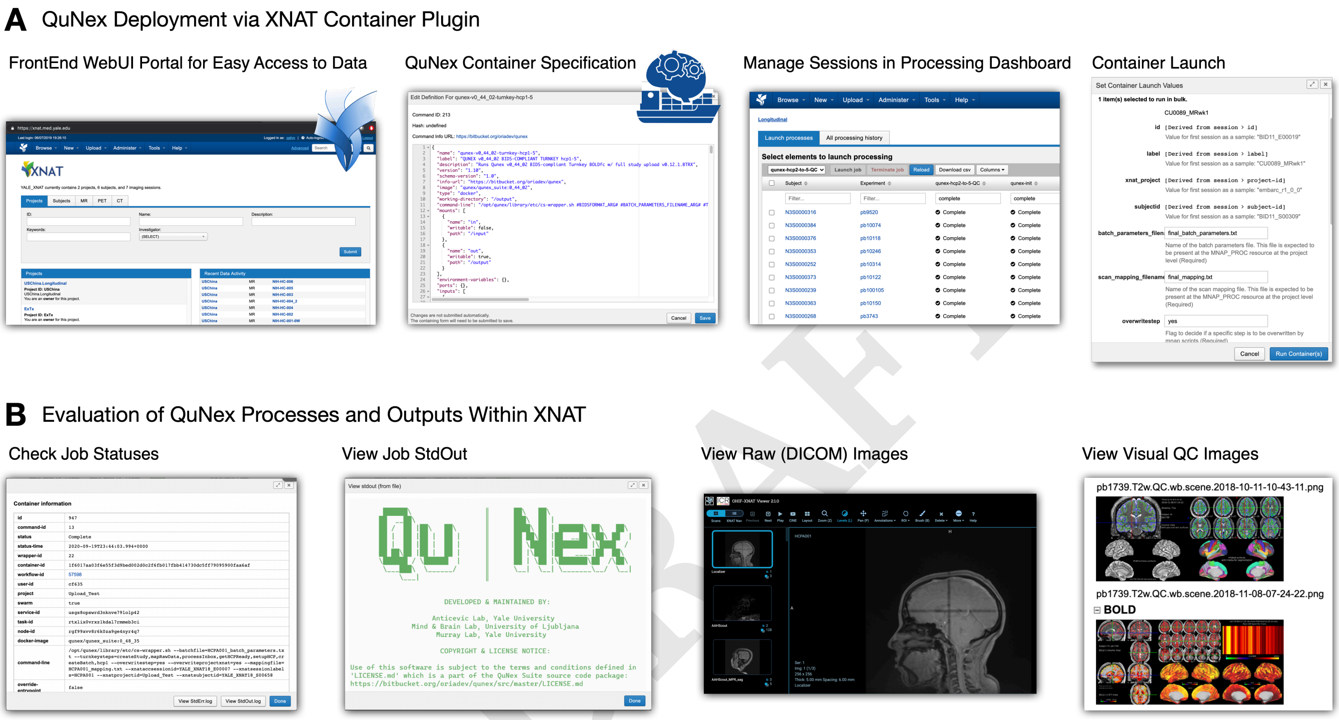


**Fig. S8. Interoperable processing and analytics with the XNAT imaging informatics platform. A)** QuNex jobs can be launched on XNAT via the container plugin. The XNAT WebUI portal allows data and studies to be easily accessed with a browser. To launch jobs, the QuNex container can be specified along with study-specific parameters in the JSON. The Processing Dashboard facilitates ability to launch containers in bulk and allows for flexible selection of sessions and container tags. Lastly, the container launch screen allows parameter selection, including specifying batch and mapping files, and turnkey steps. **B)** Intermediate and final outputs can be examined natively on XNAT to evaluate the progress of QuNex jobs. The status of executed jobs (pending, complete, error) can be easily checked, and the StdOut from each job can be accessed for information or debugging. Additionally, raw images in DICOM format and automated visual QC of processed images can be viewed within XNAT, allowing for rapid inspection of both inputs and results.

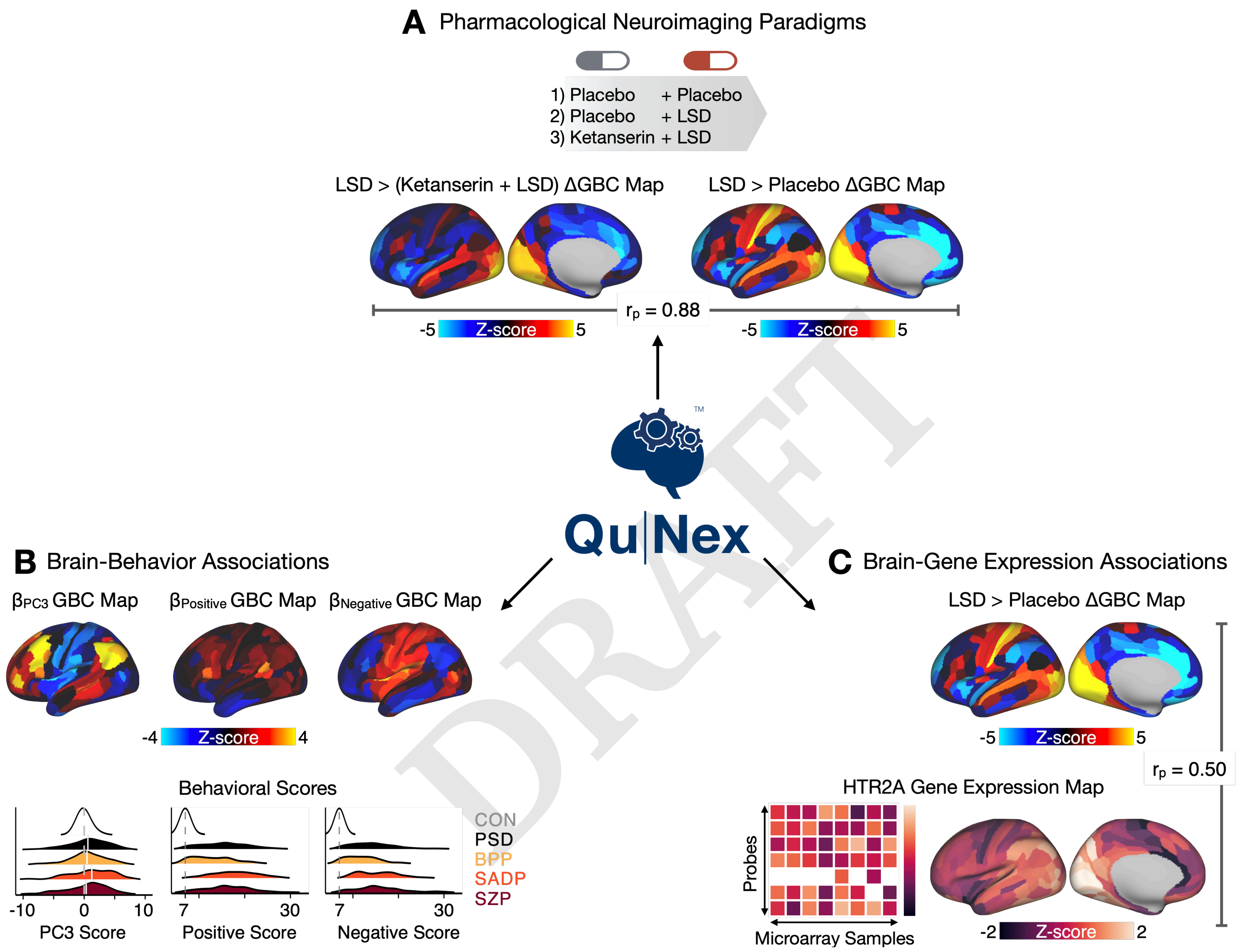


**Fig. S9. Enabling integration of independent features and surface-based analytics. A)** QuNex is capable of processing and analysis of pharmacological neuroimaging datasets designed to investigate the effect of pharmacological manipulation, such as LSD (lysergic diethylamide acid), on neuroimaging features (72). Pretreatment by ketanserin blocks the effect of LSD, resulting in a neural map that is highly similar to that of the Placebo condition. The differential global brain connectivity (ΔGBC) results between LSD and (Ket + LSD) conditions indeed show that the within-subject contrast of LSD versus (Ket + LSD) conditions is highly spatially correlated (r=0.88) with the LSD versus Placebo contrast. **B)** QuNex neural outputs can be easily quantified against behavioral phentoype data. Behavioral data, such as dimensionality-reduced symptom dimensions or composite symptom severity scores, can be mapped against neuroimaging data, such as GBC maps, across subjects in a group-level GLM. **C)** Data processed via QuNex can be seamlessly integrated with other surface-based neural features, such as cortical gene expression from the Allen Human Brain Atlas (AHBA) (85).

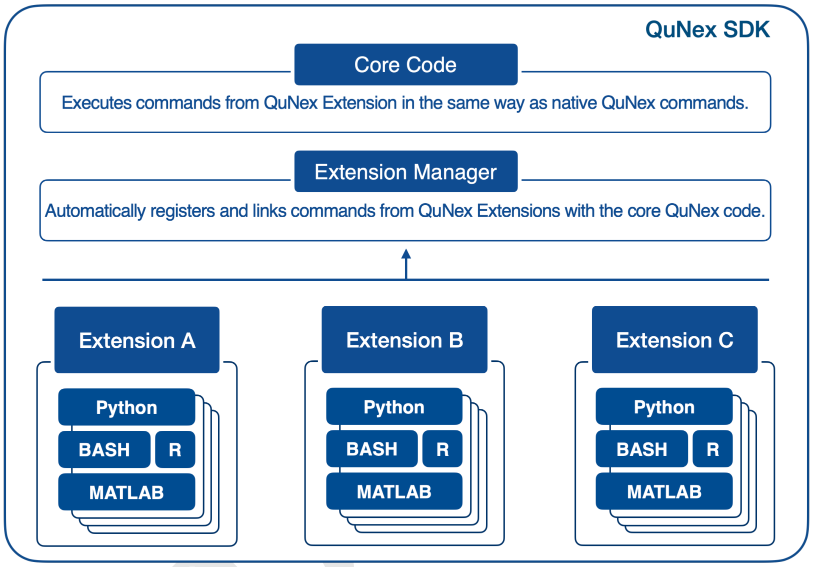


**Fig. S10. A visualization of the QuNex extensions framework.** QuNex is not open only by nature, but also by design. Through the extensions framework developers and researchers are able to design their own processing and analysis commands. These newly developed commands or extensions can facilitate all of the existing QuNex functionality (e.g. the batch turnkey engine, logging, scheduling ...). The QuNex Extensions framework guarantees organic growth and development of the platform.

| Workflow | Supported Functionality | Implementation | Existing Implementations | | | | | | | | | |
|---|---|---|---|---|---|---|---|---|---|---|---|---|
| | | | fMRIPrep | QSIPrep | HCP MPP & Workbench | UK Biobank | Nipype | micapipe | FuNP | NeuroDebian | BrainVoyager | LONI |
| Data Onboarding | BIDS import | QuNex | fMRIPrep | QSIPrep | - | - | - | micapipe | - | - | BrainVoyager | - |
| | DICOM import | dcm2niix; dcm2nii | fMRIPrep | QSIPrep | - | UK Biobank | Nibabel; dcm2nii; DCMStack | - | - | Mricron; mriconvert; heudiconv; dcm2niix; dicomnifti; Nipype | BrainVoyager | AIR |
| | Bruker import | dicomifier | - | - | - | - | BRU2NII | - | - | - | - | - |
| | NIFTI import | QuNex | fMRIPrep | QSIPrep | MPP | UK Biobank | Nibabel | - | FuNP | nifticlib; NiBabel; AFNI; caret; nipy; Nipype | BrainVoyager | AIR |
| | CIFTI | QuNex; HCP | fMRIPrep | QSIPrep | MPP | UK Biobank | Nibabel | HCP | HCP | NiBabel; HCP; caret | BrainVoyager | - |
| Preprocessing | T1w registration | FSL; ANTs | FSL; ANTs | ANTs | FSL | FSL | FSL; AFNI; ANTS; BrainSuite; Nifty; SPM | FSL; AFNI; Elastix | FSL; AFNI | FSL; AFNI; SPM; HCP | BrainVoyager | FSL; AFNI; SPM; BrainSuite |
| | T1 segmentation | FreeSurfer | FreeSurfer | - | FreeSurfer | FreeSurfer | FreeSurfer; BrainSuite; Nifty | FreeSurfer | FSL | FreeSurfer; itksnap | BrainVoyager | FSL; AFNI; SPM; BrainSuite |
| | T2w registration | FSL; ANTs | FSL; ANTs | - | Freesurfer | FSL | FSL; AFNI; ANTs; Nifty | - | - | FSL; AFNI; SPM; HCP | BrainVoyager | FSL; AFNI; SPM; BrainSuite |
| | BOLD | FSL; FreeSurfer; HCP | ANTs; FSL; FreeSurfer | - | MPP | FSL | FSL; AFNI; Nifty; SPM; Vistasoft; HCP | AFNI; FSL; HCP | FSL; HCP | FSL; AFNI; Nifty; SPM; HCP; NiPy; Nitime | BrainVoyager | FSL; AFNI; SPM; BrainSuite |
| | DWI | HCP; FSL | - | ANTs; FSL; MRtrix3; DSL Studio; DIPY | MPP; FSL | FSL | VistaSoft; MRtrix3; Camino; DIPY; HCP; DTITK; Diffusion Toolkit; FSL; BrainSuite | MRtrix3 | - | FSL; HCP | BrainVoyager | FSL; Diffusion Toolkit; BrainSuite |
| | PET | Staged for development | - | - | - | - | PETPVC | - | - | - | BrainVoyager | - |
| | EEG | Under development | - | - | - | - | MNE | - | - | Openmeeg | BrainVoyager | - |
| | Distortion correction | FSL; ANTs | FSL; ANTs | FSL; ANTs | FSL; HCP | FSL; HCP | FSL; AFNI; ANTs; BrainSuite; Nifty; SPM; HCP; VistaSoft; MRtrix3; Camino; DIPY; DTITK; Diffusion Toolkit; BrainSuite | AFNI; FSL; HCP | AFNI | AFNI; FSL; HCP; Nipype | BrainVoyager | FSL; AFNI; SPM; BrainSuite |
| | Motion correction | FSL; ANTs | FSL | FSL; QSIPrep | FSL | FSL | FSL; AFNI; ANTs; BrainSuite; Nifty; SPM; HCP; VistaSoft; MRtrix3; Camino; DIPY; HCP; DTITK; Diffusion Toolkit; FSL; BrainSuite | AFNI; FSL; HCP | AFNI | AFNI; FSL; HCP; Nipype; SPM | BrainVoyager | FSL; AFNI; SPM; BrainSuite |
| | Longitudinal preprocessing | FreeSurfer | - | - | - | - | FreeSurfer | - | - | - | - | - |
| Denoising | Component-based nuisance removal | ICA-FIX; tICA | ICA-AROMA; CompCor | - | ICA-FIX; tICA | ICA-FIX | ICA-AROMA; CompCor | ICA-FIX | ICA-FIX | ICA-AROMA | BrainVoyager | ICA-AROMA |
| | Motion scrubbing | QuNex | AFNI; fMRIPrep | - | - | - | AFNI | FSL | FSL | FSL; AFNI | BrainVoyager | FSL; AFNI |
| | Nuisance regression | QuNex | FSL; fMRIPrep | - | - | FSL | FSL; AFNI; Nifty; SPM; HCP; NiPy; Nitime | FSL | - | FSL; AFNI; Nifty; SPM; HCP; NiPy; Nitime | BrainVoyager | AFNI; FSL; HCP |
| | Temporal filtering | QuNex | FSL | - | - | FSL | FSL; AFNI; Nifty; SPM; HCP | FSL | AFNI | FSL; AFNI; Nifty; SPM; HCP; NiPy; Nitime | BrainVoyager | AFNI; FSL; HCP |
| | Spatial smoothing | QuNex | AFNI | - | Workbench | FSL | FSL; AFNI; Nifty; SPM; HCP; NiPy; Nitime | FSL | AFNI | FSL; AFNI; Nifty; SPM; HCP; NiPy; Nitime | BrainVoyager | AFNI; FSL; HCP |
| Quality Control | Visual DICOM QC | QuNex | - | - | - | - | - | - | - | - | - | LONI |
| | Visual T1w QC | QuNex | fMRIPrep | - | MPP | - | - | micapipe | - | HCP | - | LONI |
| | Visual T2w QC | QuNex | fMRIPrep | - | MPP | - | - | - | - | HCP | - | LONI |
| | Visual Myelin QC | QuNex | fMRIPrep | - | MPP | - | - | micapipe | - | HCP | - | LONI |
| | Visual DWI QC | QuNex | fMRIPrep | QSIPrep | MPP | - | - | micapipe | - | HCP | - | LONI |
| | Visual BOLD QC | QuNex | fMRIPrep | - | MPP | - | - | micapipe | - | HCP | - | LONI |
| | Motion plots | QuNex | fMRIPrep | - | MPP | - | - | micapipe | - | HCP | - | LONI |
| | Automated QC | Under development | - | - | - | UK Biobank | - | - | - | - | - | LONI |
| Feature Generation | Parcellating timeseries | QuNex; Workbench | - | - | Workbench | FSL | FSL; AFNI; SPM; HCP | AFNI; FSL; HCP | - | FSL; AFNI; SPM; HCP; VoxBo | BrainVoyager | FSL; AFNI; SPM |
| | Functional connectivity | QuNex; Workbench | - | - | Workbench | FSL | FSL; AFNI; SPM; HCP | micapipe | - | FSL; AFNI; SPM; HCP | BrainVoyager | FSL; AFNI; SPM |
| | Extracting task activation maps | QuNex | - | - | - | FSL | FSL; AFNI; SPM | FSL | - | FSL; AFNI; SPM | BrainVoyager | FSL; AFNI; SPM |
| | Tractography | FSL | - | ANTs; FSL; MRtrix3; DSL Studio; DIPY | FSL | FSL | VistaSoft; MRtrix3; Camino; DIPY; HCP; DTITK; Diffusion Toolkit; FSL; BrainSuite | MRtrix3; ANTs | - | FSL | BrainVoyager | FSL; Diffusion Toolkit; BrainSuite |
| | Diffusion measures | FSL | - | ANTs; FSL; MRtrix3; DSL Studio; DIPY | FSL | FSL | VistaSoft; MRtrix3; Camino; DIPY; HCP; DTITK; Diffusion Toolkit; FSL; BrainSuite | MRtrix3; ANTs | - | FSL | BrainVoyager | FSL; BrainSuite; Diffusion Toolkit |
| | Structural measures | FreeSurfer | FreeSurfer | - | FreeSurfer | FreeSurfer | FreeSurfer; BrainSuite; ANTs; Nifty | FreeSurfer | - | FreeSurfer | BrainVoyager | FreeSurfer |
| Group Level Analytics | Non-parametric permutation inference | FSL | - | - | - | FSL | FSL; AFNI; SPM | - | - | FSL; AFNI; SPM | BrainVoyager | FSL; AFNI; SPM |
| | Spatial statistics | FSL | - | - | - | FSL | FSL; AFNI; SPM | - | - | FSL; AFNI; SPM | BrainVoyager | FSL; AFNI; SPM |
| | Joint inference tests | FSL | - | - | - | FSL | FSL | - | - | FSL | BrainVoyager | FSL |
| | Multi-modality tests | FSL | - | - | - | FSL | FSL | - | - | FSL | BrainVoyager | FSL |
| Species Support | Macaque | FSL; HCP | - | - | MPP; Workbench | - | FSL | - | - | FSL | BrainVoyager | - |
| | Mouse | Under development | - | - | Workbench | - | semTools | - | - | - | - | - |
| Deployment Options | Turnkey functionality | QuNex | Yes | - | - | Yes | Yes | - | Yes | - | BrainVoyager | Yes |
| | Container | QuNex | Yes | - | - | - | - | Yes | - | Yes | BrainVoyager | - |
| | Native scheduler support | QuNex | - | - | - | - | Yes | - | - | - | - | Yes |
| | Cloud deployment | QuNex | Yes | - | - | - | Yes | - | - | Yes | - | Yes |
| | API | Staged for development | - | - | - | - | Yes | - | - | Yes | BrainVoyager | Yes |
| Development | Interoperability of tools | Yes | Yes | Yes | - | Yes | Yes | Yes | Yes | Yes | Yes | Yes |
| | Open source code | Yes | Yes | Yes | Yes | Yes | Yes | Yes | Yes | Yes | - | Yes |
| | Software Development Kit | Yes | - | - | - | - | Yes | - | - | Yes | - | Yes |
| | Extensions framework | Yes | - | - | - | - | Yes | - | - | - | - | Yes |
| User Support | Online documentation | Yes | Yes | Yes | Yes | Yes | Yes | Yes | - | Yes | Yes | Yes |
| | Community forum | Yes | Yes | Yes | Yes | - | - | Yes | - | Yes | Yes | Yes |
| | Active maintenance | Yes | Yes | Yes | Yes | Yes | Yes | Yes | Yes | Yes | Yes | - |

**Fig. S11. Features included in QuNex and comparisons to other neuroimaging software.** Here we list the implmentations of tools for supported QuNex functionalities. We also list existing implementations in other currently available neuroimaging pipelines/environments, including fMRIPrep (10), QSIPrep (11), HCP (9), UK Biobank (38), nipype (17), micapipe (39), FuNP (41), NeuroDebian (44), BrainVoyager (40), and LONI (45).
